# KSR1- and ERK-dependent Translational Regulation of the Epithelial-to-Mesenchymal Transition

**DOI:** 10.1101/2021.01.18.427224

**Authors:** Chaitra Rao, Danielle E. Frodyma, Siddesh Southekal, Robert A. Svoboda, Adrian R. Black, Chittibabu Guda, Tomohiro Mizutani, Hans Clevers, Keith R. Johnson, Kurt W. Fisher, Robert E. Lewis

## Abstract

The epithelial-to-mesenchymal transition (EMT) is considered a transcriptional process that induces a switch in cells from a polarized state to a migratory phenotype. Here we show that KSR1 and ERK promote EMT through the preferential translation of Epithelial-Stromal Interaction 1 (EPSTI1), which is required to induce the switch from E-to N-cadherin and coordinate migratory and invasive behavior. EPSTI1 is overexpressed in human colorectal cancer (CRC) cells. Disruption of KSR1 or EPSTI1 significantly impairs cell migration and invasion *in* vitro, and reverses EMT, in part, by decreasing the expression of N-cadherin and the transcriptional repressors of E-cadherin expression, ZEB1 and Slug. In CRC cells lacking KSR1, ectopic EPSTI1 expression restored the E-to N-cadherin switch, migration, invasion, and anchorage-independent growth. KSR1-dependent induction of EMT via selective translation of mRNAs reveals its underappreciated role in remodeling the translational landscape of CRC cells to promote their migratory and invasive behavior.

## Introduction

Molecular scaffolds affect the intensity and duration of signaling pathways by coordinating a discrete set of effectors at defined subcellular locations to regulate multiple cell fates (1, 2). Kinase Suppressor of Ras 1 (KSR1) serves as a scaffold for Raf, MEK, and ERK enabling the efficient transmission of signals within the mitogen activated protein kinase (MAPK) cascade (3, 4). Although KSR1 is dispensable for normal development, it is necessary for oncogenic Ras-induced tumorigenesis including colorectal cancer cells (3-7), suggesting that KSR1 may modulate aberrant signals that redirect the function of effectors typically involved in normal cellular homeostasis. Activating Ras mutations are present in over 40% of colorectal cancers (CRC), and associated with advanced disease and decreased overall survival (8, 9). Activated Ras, a critical driver of both tumor growth and survival, is an alluring therapeutic target, yet targeting the majority of oncogenic Ras alleles is still a work in progress. Raf/MEK/ERK signaling can phenocopy Ras signaling essential for CRC growth and survival (10, 11). Therefore, understanding the effectors that transmit signals emanating from oncogenic Ras is a valuable step in detecting and targeting the pathways critical to tumor cell function and their adaptation to therapy.

Oncogene-driven signaling pathways promote protein translation that enables expression of a subset of mRNAs to promote growth, invasion, and metastasis (12-15). Tumor cells have an increased dependence on cap-dependent translation, unlike their normal complements (14, 16). Eukaryotic Translation Initiation Factor 4E (eIF4E) is a rate-limiting factor for oncogenic transformation, with reductions of as little as 40% being sufficient to block tumorigenesis (14). eIF4E function is regulated by association of 4E-binding proteins (4EBPs). Importantly, disruption of KSR1 or ERK inhibition leads to dephosphorylation and activation of 4EBP1, indicating that the function of KSR1 as an ERK scaffold is key to the aberrant regulation of protein translation (17). This tumor-specific, KSR1-dependent regulation of protein translation of a subset of genes was predicted to selectively promote survival of CRC cells but not normal colon epithelia (17, 18).

Almost all CRC originates from epithelial cells lining the colon or rectum of the gastrointestinal tract, but in order to invade to the surrounding tissue, cancer cells lose cell adhesiveness to acquire motility and become invasive, characterized by the epithelial-to-mesenchymal transition (EMT), which is central to tumor pathogenesis (19-22). EMT involves a complex cellular process during which epithelial cells lose polarity, cell-cell contacts and acquire mesenchymal characteristics. While EMT is crucial for cell plasticity during embryonic development, trans differentiation and wound healing, when aberrantly activated EMT has deleterious effects, which facilitate motility and invasion of cancer cells (20-23). EMT has been shown to be controlled by transcription-dependent mechanisms, especially through repression of genes that are hallmarks of epithelial phenotype such as E-cadherin. Loss of E-cadherin at the membrane has been associated with carcinoma progression and EMT (21, 24-26). E-cadherin function is transcriptionally repressed through the action of EMT transcription factors (TFs), including Snail-family proteins (*Snail1, Slug*), zinc finger E-box binding homeobox 1 and 2 (*ZEB1* and *ZEB2*) and twist-related protein (*Twist*) (23, 27). Transcriptional control of E-cadherin is unlikely to be sole determinant of EMT, invasion and metastasis. Inappropriate induction of non-epithelial cadherins, such as N-cadherin by epithelial cells are known to play a fundamental role during initiation of metastasis (28-34). N-cadherin disassembles adherent junction complexes, disrupting the intercellular cohesion and reorienting the migration of cells, away from the direction of cell-cell contact (28, 35). Upregulation of N-cadherin expression promotes motility and invasion (28-30, 36). Thus, central to the process of EMT is the coordinated loss of E-cadherin expression and the upregulation of N-cadherin gene expression, termed cadherin switching (34, 37-40).

Previous studies have demonstrated transcriptional regulation of EMT through oncogenic Ras or its downstream effector signaling pathways via the activation of EMT-TFs (41-47). Oncogenic Ras itself activates EMT-TF *Slug* to induce EMT in skin and colon cancer cells (45, 46). Enhanced activity of ERK2 but not ERK1, has been linked with Ras-dependent regulation of EMT (41, 42). Several studies have also described an alternative program wherein cells lose their epithelial phenotype, via post-transcriptional modifications rather than transcriptional repression involving translational regulation or protein internalization (48-50). Expression profiling of polysome-bound mRNA to assess translational efficiency identified over thirty genes that were translationally regulated upon Ras and *TGFβ* inducing EMT (48, 50). Functional characterization of the resultant proteins should reveal preferentially translated mRNAs essential to invasion and metastasis.

*EPSTI1* was identified as a stromal fibroblast induced gene upon co-cultures of breast cancer cells with stomal fibroblasts (51). EPSTI1 is expressed at low levels in normal breast and colon tissue but aberrantly expressed in breast tumor tissue (51). EPSTI1 promotes cell invasion and malignant growth of primary breast tumor cells (52, 53). We performed polysome profiling in CRC cells and found that KSR1- and ERK induces of EPSTI1 protein translation. EPSTI1 is both necessary and sufficient for coordinating the up-regulation of N-cadherin with the downregulation of E-cadherin to stimulate cell motility and invasion in colon cancer cells. These data demonstrate that ERK-regulated regulation of protein translation is an essential contributor to EMT and reveal a novel effector of the cadherin switch whose characterization should yield novel insights into the mechanisms controlling the migratory and invasive behavior of cells.

## Results

### Genome wide polysome profiling reveals translational regulation of EPSTI1 by KSR1

ERK signaling regulates global and selective mRNA translation through RSK1/2-dependent modification of cap-dependent translation (17, 54). Phosphorylation of cap binding protein 4E-BP1 releases eIF4E to promote translation and the abundance of eIF4E is a rate-limiting factor for oncogenic Ras- and Myc-driven transformation (14). We showed previously that KSR1 maximizes ERK activation in the setting of oncogenic Ras (55), which is required for increased Myc translation via dephosphorylation of 4E-BP1, supporting CRC cell growth (17). These observations imply that the ERK scaffold function of KSR1 alters the translational landscape in CRC cells to support their survival.

To determine the effect of KSR1 on translatomes in colon cancer cells, we performed genome-wide polysome profiling (56). We stably expressed short hairpin RNA (shRNA) constructs targeting KSR1 (KSR1 RNAi) or a non-targeting control in two K-Ras mutant CRC cell lines, HCT116 and HCT15 **(Fig. 1D, top panels)**. We isolated and quantified both total mRNA and efficiently translated mRNAs (associated with ≥ 3 ribosomes) using RNA sequencing (**Fig. 1A**). We used Anota2seq (57) to calculate translation efficiency (TE) by comparing the differences in efficiently translated mRNAs to the total transcript of each mRNA and observed that a significant number of mRNAs ([selDeltaTP ≥ log (1.2)] and selDeltaPT ≥ log (1.2)] and p value < 0.05) showed either reduced TE or upregulated TE upon KSR1 disruption (**Fig. 1B-C, Supplementary Table 1**) in both HCT116 and HCT15 cells. Gene Set Enrichment Analysis (GSEA) of significantly enriched genes in HCT116 and HCT15, identified 11 mRNAs (**Fig. 1B, supplementary Fig. 1A**) in the gene set titled “Hallmark EMT signature”, “Jechlinger EMT Up”, and Gotzmann EMT up” (58), that had significantly decreased translation upon KSR1 disruption (**Supplementary Table 2**). Among the genes with decreased translation, *EPSTI1* was one of the highly significant mRNAs. We sought to determine the functional relevance of KSR1-dependent induction of EPSTI1 to EMT in colon cancer cells.

**Figure 1.**
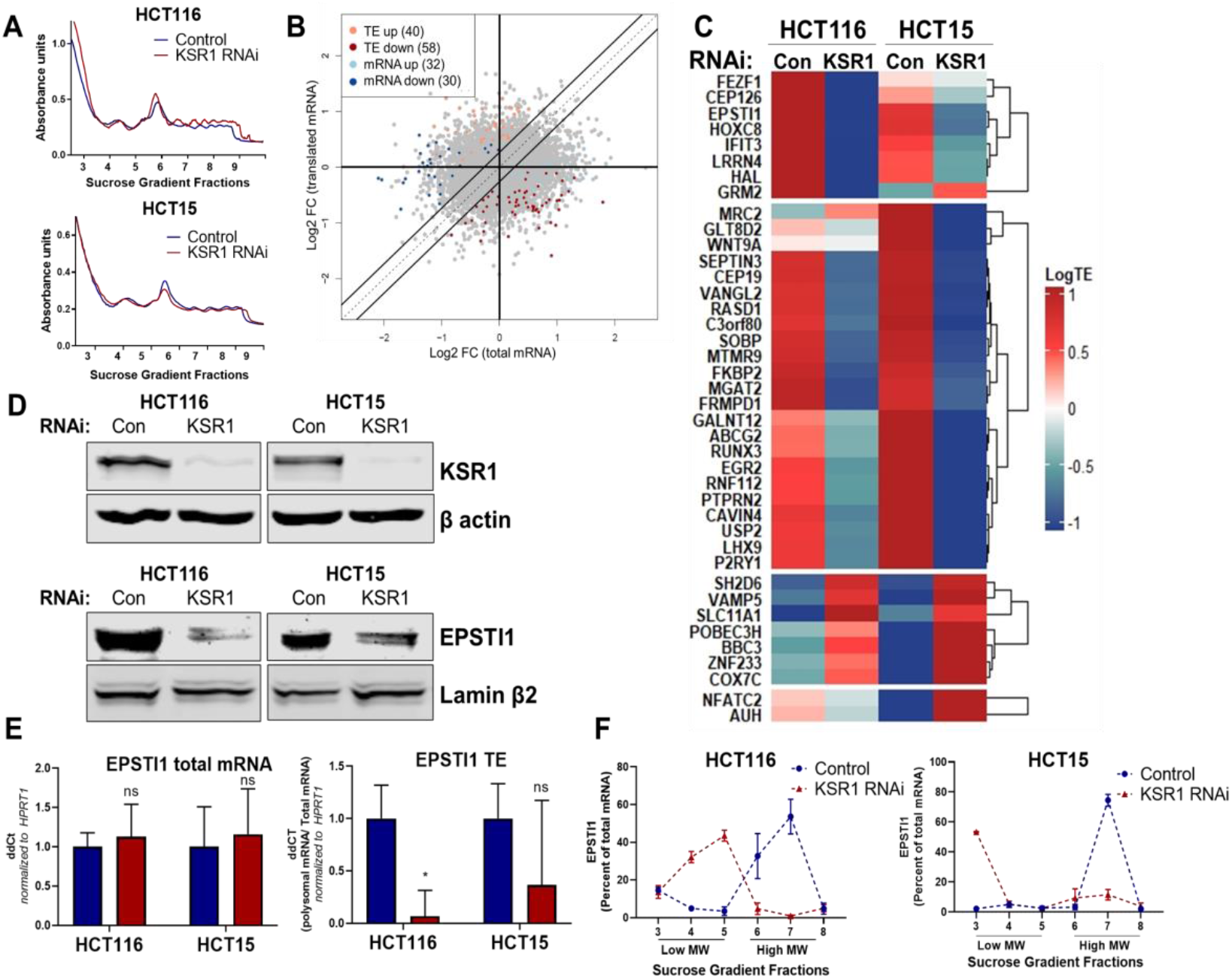
EPSTI1 translation is regulated by KSR1. **(A)** Representative polysome profiles from control and KSR1 knockdown (KSR1 RNAi) HCT116 and HCT15 cells. Sucrose gradient fractions 3-5 denote the low molecular weight complexes and the fractions 6-9 are the high molecular weight polysomes. **(B)** Scatter plot of polysome-associated mRNA to total mRNA log2 fold-changes upon KSR1 knockdown in HCT116 and HCT15 with RNA-seq. The statistically significant genes in the absence of KSR1 are classified into four groups with a fold change (|log_2_FC|) > 1.2 and p-value < 0.05. The number of mRNAs with a change in TE (orange and red) are indicated (n=3 for each condition). TE, translational efficiency. **(C)** Heatmap of TE changes for the top 40 RNAs control and KSR1 knockdown (KSR1 RNAi) HCT116 and HCT15 cells (n=3 for each condition). **(D)** Western blot analysis of KSR1 and EPSTI1 following KSR1 knockdown in HCT116 and HCT15 cells. **(E)** RT-qPCR analysis of EPSTI1 mRNA from total RNA and polysomal RNA (fractions number 6-8) in control and KSR1 knockdown HCT116 and HCT15 cells, the TE was calculated as the ratio of polysomal mRNA to the total mRNA (n=3; *, P <0.05). **(F)** RT-qPCR analysis of *EPSTI1* mRNA levels isolated from sucrose gradient fractions of the control and KSR1 knockdown HCT116 and HCT15 cells. Fractions 3-5 (low MW) and 6-8 (high MW) are plotted for the control and KSR1 knockdown state with values corresponding to the percentage of total mRNA across these fractions n=3. Experiments shown in (A - F) are representative of three independent experiments.

To confirm that EPSTI1 translation is KSR1-dependent, we observed that, EPSTI1 protein expression was decreased with the knockdown of KSR1 in HCT116 and HCT15 cells (**Fig. 1D**), while the total mRNA transcript was unchanged upon KSR1 disruption (**Fig. 1E**, left panel). *EPSTI1* TE was markedly decreased upon KSR1 depletion (**Fig. 1E**, right). RT-qPCR analysis of sucrose-gradient fractions of monosome mRNA and polysome RNA distribution confirmed that *EPSTI1* mRNA shifted from actively translating high molecular weight (MW) polysome fractions to low MW fractions in KSR1 knockdown cells (**Fig. 1F**). In contrast, *HPRT1* mRNA was insensitive to KSR1 knockdown in HCT116 and HCT15 cells, and qPCR analysis of *HPRT1 mRNA* isolated from sucrose gradient fractions of control and KSR1 knockdown cells showed no significant shift between the low MW and the high MW fractions **(Supplementary Fig. 1C)**. These data show EPSTI1 translation is induced by KSR1.

### KSR1/ERK signaling regulates EPSTI1 expression in colon cancer cells

To confirm our observations in KSR1 knockdown cells, we tested the effect of CRISPR/Cas9-mediated targeting of KSR1 on EPSTI1 in CRC cell lines. EPSTI1 protein expression was decreased upon KSR1 depletion in HCT116 and HCT15 cells and EPSTI1 expression was restored in knockout cells upon expression of a KSR1 transgene (+ KSR1) (**Fig. 2A**). Similar to inhibition of KSR1, treatment with ERK inhibitor SCH772984 (59) suppressed EPSTI1 protein expression in both CRC cell line HCT116 and tumorigenic patient derived colon organoid engineered with deletion of APC, p53, SMAD4 and K-Ras^G12D^ mutation (PDO-11 AKPS) **(Fig. 2B)** (60). While the total protein was reduced upon ERK inhibition in HCT116, the *EPSTI1* transcript levels were not altered significantly by SCH772984 treatment **(Fig. 2C)**.

**Figure 2.**
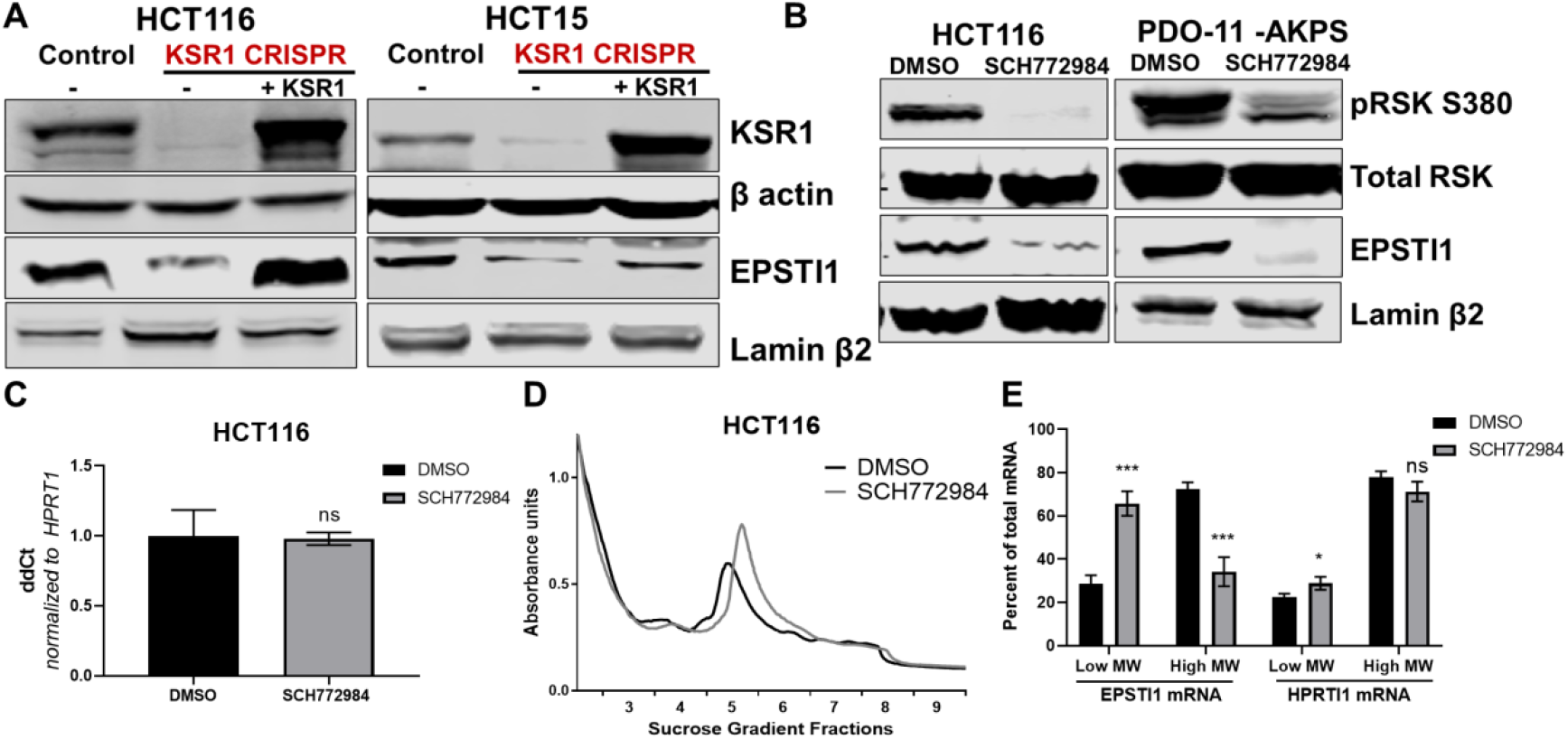
KSR1 or ERK inhibition suppresses EPSTI1 protein expression in cell lines and organoids. **(A)** Cell lysates prepared from control, KSR1 CRISPR-targeted (KSR1 CRISPR) and CRISPR-targeted HCT116 and HCT15 cells expressing KSR1 (KSR1 CRISPR + KSR1) analyzed for EPSTI1 protein expression by Western blotting. **(B)** Western blot of the indicated proteins in HCT116 (left) and AKPS quadruple mutant organoids (right) treated with DMSO or 1 µM of SCH772984 for 48 hours. **(C)** RT-qPCR analysis of *EPSTI1* mRNA from total RNA in HCT116 cells treated with either DMSO or ERK1/2 selective inhibitor, SCH772984 (n=3; ns, non-significant). **(D)** Representative polysome profiles from HCT116 cells treated DMSO or 1 µM of ERK1/2 selective inhibitor, SCH772984. **(E)** RT-qPCR analysis of *EPSTI1* and *HPRT1* mRNA levels from LMW (fractions 3-5) and HMW (fractions 6-8) of the DMSO control and SCH772984-treated HCT116 cells (n=3; *, P<0.05; ***, P<0.001). All values displayed as mean ± S.D. Experiments shown in (A-E) are representative of three independent experiments.

We performed polysome profiling in HCT116 cells, either treated with DMSO or ERK inhibitor, SCH772984 and we isolated mRNA from low MW monosome (fractions 3-5) and high MW polysome (fractions 6-8) fractions (**Fig. 2D**). RT-qPCR demonstrated that *EPSTI1* mRNA shifted from high MW fractions to the low MW fractions upon ERK inhibition **(Fig. 2E)**. The distribution of mRNA for *HPRT1* within the same profile was not altered by SCH772984 treatment (**Fig. 2E**). These data indicate that KSR1-dependent ERK signaling is a critical regulator of EPSTI1 protein translation in colon cells and organoids.

### EPSTI1 is required for anchorage-independent growth in colon cancer cells

KSR1 disruption inhibits HCT116 cell anchorage-independent growth *in vitro* and tumor formation *in vivo* (6). Similarly, disruption of KSR1 by CRISPR/Cas9-mediated targeting decreased HCT116 and HCT15 cell viability under anchorage-independent conditions on simulated by poly-(HEMA) coating (**Fig. 3A**). KSR1 transgene expression restored cell viability in HCT116 and HCT15 cells lacking KSR1 (KSR1 CRISPR + KSR1) (**Fig. 3A**). EPSTI1 protein is aberrantly expressed in colon cancer cell lines HCT116 and HCT15, while the expression is detected weakly in non-transformed human colon epithelial cells (HCECs) (**Fig. 3B**). EPSTI1 protein expression is also markedly higher in AKPS organoids than normal colon organoids (**Fig. 3B**).

**Figure 3.**
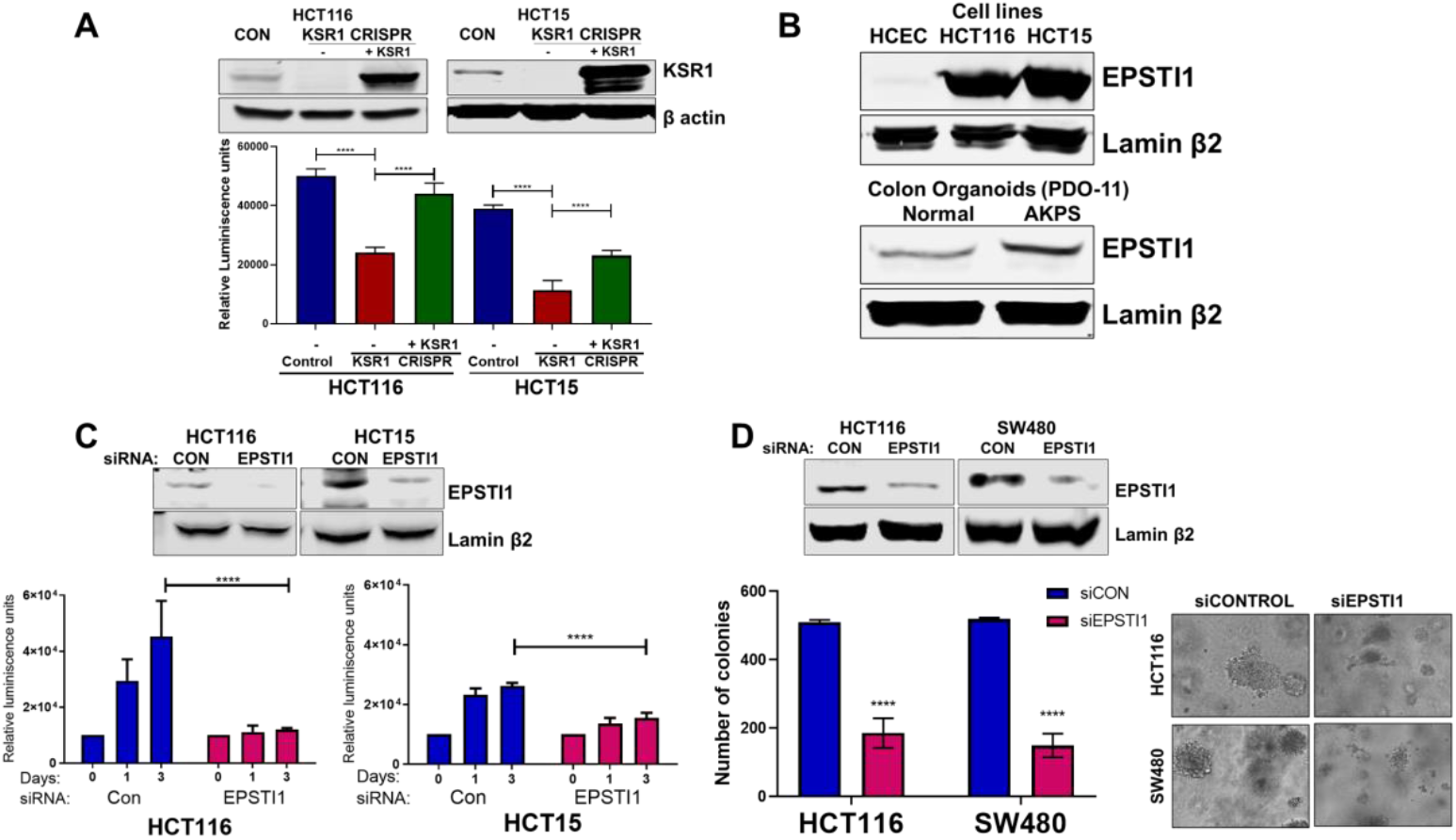
EPSTI1 is overexpressed in cancer cell lines and organoids and promotes anchorage-independent growth. **(A)** Anchorage-independent cell viability was analyzed in HCT116 and HCT15 cells plated on poly-(HEMA)-coated plates was measured using CellTiter-Glo following CRISPR-targeting (KSR1 CRISPR) and re-expressing KSR1 (KSR1 CRISPR + KSR1) in the CRISPR-targeted cells. The data are shown as relative luminescence units mean ± SD, n=6. Matched results were analyzed for statistical significance one-way ANOVA followed by t-test. (Upper panels) Western blot showing the expression of KSR1 in control, KSR1 knockout and KSR1-knockout cells expressing a KSR1 transgene (+ KSR1). **(B)** Western blot analysis of EPSTI1 protein expression was assessed in HCECs, HCT116, HCT15, normal human colon organoids, and transformed AKPS colon organoids. **(C)** Viability of HCT116 and HCT15 cells measured using CellTiter-Glo following siRNA knockdown of EPSTI1 that were plated on poly-(HEMA)-coated plates to simulate anchorage-independent conditions. Cell viability was measured immediately after plating and 0, 1 and 3 days after plating (n=6). The data are shown as mean luminescence units ± SD. Matched results were analyzed for statistical significance by t-test. (Top) Western blot confirming the knockdown of EPSTI1 in HCT116 and SW480 at Day 3. **(D)** (Left) Quantification of the colonies formed in HCT116 and SW480 cells following RNAi knockdown using non-targeting control (siCON) or EPSTI1 (siEPSTI1) after plating on soft agar. (Right) Representative photomicrographs of colonies for each sample. The data are illustrated as the number of colonies present after two weeks, mean ± SD, n=6. Paired results were analyzed for statistical significance using Student’s *t* test. (Top) Western blot confirming the knockdown of EPSTI1 in HCT116 and SW480 cells. ****, P < 0.0001

To determine the regulation of EPSTI1 in human colon tumor maintenance, we performed siRNA knockdown of EPSTI1 in HCT116 and HCT15 cells. EPSTI1 disruption suppressed viability on poly-(HEMA) coated by 40% in HCT15 cells, and over 70%, in HCT116 cells (**Fig. 3C**). EPSTI1 knockdown reduced colony formation in soft agar by 63% in HCT116 cells and 71% in SW480 cells (**Fig. 3D**). These observations show that KSR1-dependent translation of ESPTI1 is required for colon tumor cell transformation.

### KSR1 or EPSTI1 disruption decreases cell mobility in CRC cells

Considering the suggested role of EPSTI1 in promoting EMT-like phenotypes (51, 52), we sought to evaluate the biological role of EPSTI1 in colon cancer cells. Time-lapse images of control and EPSTI1 knockdown in HCT116 cell motility in a scratch wound was analyzed by measuring the relative wound density (61) over 72 hours (**Fig. 4A, bottom**). Motility was also assessed in control, CRISPR-targeted (KSR1 CRISPR), and CRIPSR-targeted HCT116 cells expressing KSR1 (KSR1 CRISPR + KSR1) (**Fig. 4A, top**). Cells lacking either EPSTI1 or KSR1 were approximately 20% less motile compared to control cells. Reintroduction of KSR1 expression in CRISPR-targeted HCT116 cells restored motility comparable to the control cells (**Fig. 4A, top**).

**Figure 4.**
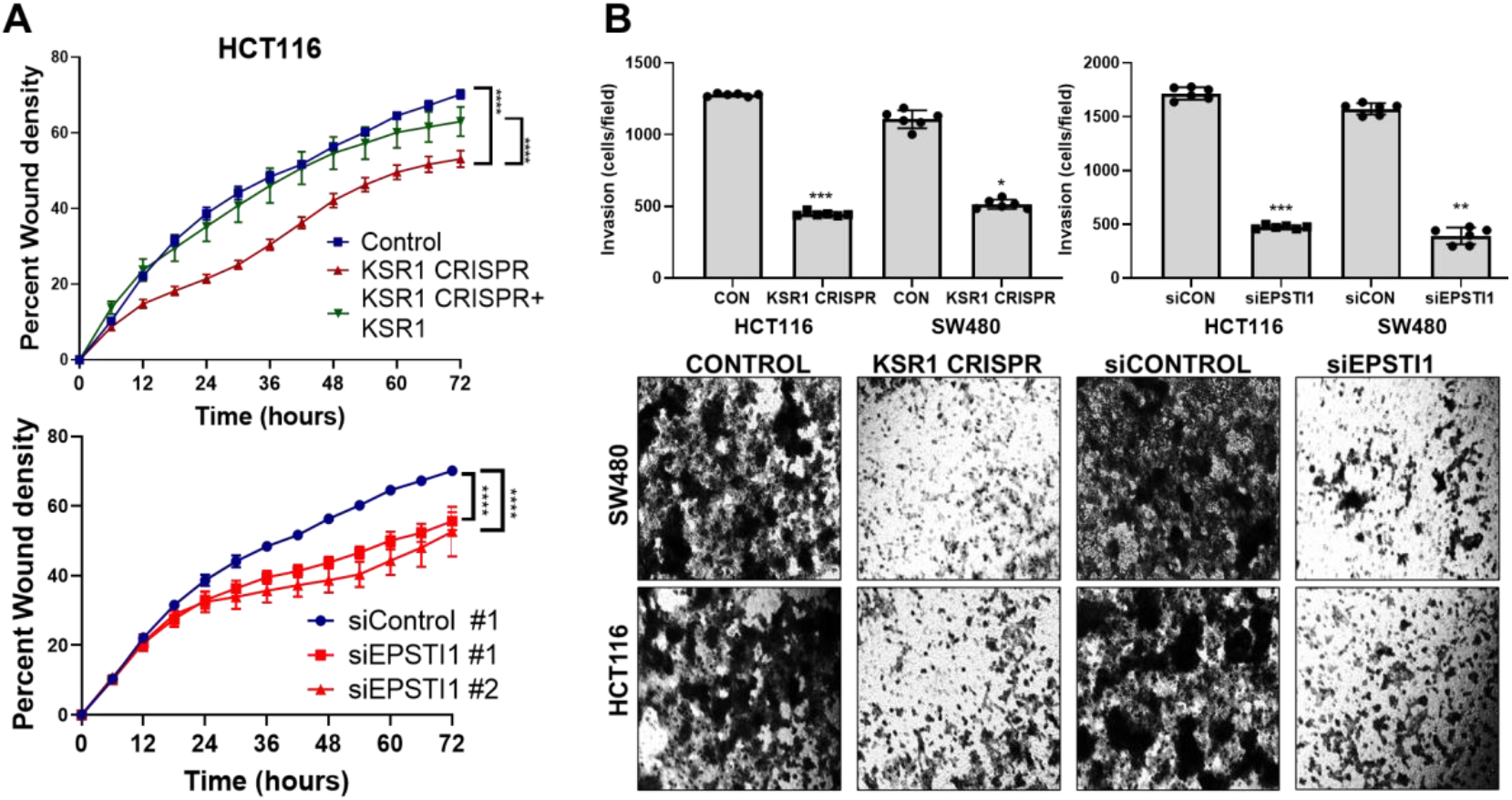
KSR1 and EPSTI1 promote migration and invasion in CRC cells. **(A)** Control, CRISPR-targeted (KSR1 CRISPR) and CRISPR-targeted HCT116 cells expressing KSR1 (KSR1 CRISPR + KSR1) (upper) and control or EPSTI1 knockdown HCT116 cells (lower) were evaluated in a 96-well IncuCyte scratch wound assay. The graph represents the time kinetics of percent wound density, calculated by IncuCyte ZOOM software, shown as mean ± SD, n=12 ****, P < 0.0001. Matched results were analyzed for statistical significance using one-way ANOVA with Dunnett’s posttest for multiple comparisons. **(B)** (Upper panels) Control, KSR1 knockout (KSR1 CRISPR) and EPSTI1 knockdown (siEPSTI1) were subjected to Transwell migration assay through Matrigel^®^ for 24 hours using 10% FBS as chemoattractant. The number of invaded cells per field were counted. Data are the mean ± SD (n=6); *, P < 0.1; **, P < 0.01; ***, P < 0.001. (Lower panels) Representative images of Giemsa-stained cells 24 hours after invasion through Matrigel^®^.

EPSTI1 knockdown HCT116 and SW480 cells were subjected to Transwell invasion assays. EPSTI1 RNAi suppresses cell invasion through Matrigel^®^ by 72% in HCT116 and by 75% in SW480. (**Fig. 4B, top right and bottom**). Since KSR1 is required for EPSTI1 translation, we determined the functional contribution of KSR1 in regulating cell invasion. KSR1 depletion suppressed invasion by 64% in HCT116 and by 53% SW480 cells (**Fig. 4B, top left and bottom**). Overall, these results suggest the KSR1-dependent EPSTI1 signaling contributes to cell migration and invasion in CRC cells.

### KSR1 or EPSTI1 disruption causes cadherin switching in CRC cells

To understand the underlying mechanism by which KSR1 and EPSTI1 promote motility and invasion in CRC cells, we evaluated their contribution to the expression of critical determinants of EMT that modulate cell adhesion, E- and N-cadherins and EMT-TFs. Compared to the non-targeting control, KSR1 disruption in HCT116, HCT15 and SW480 cells had elevated levels of E-cadherin, along with a coincident decrease in EMT-TF Slug **(Fig. 5A)**. Expression of Vimentin, and Snail1 was not changed in HCT116 cells **(Supplementary Fig. 2A)**. Upon knockdown of EPSTI1 with either of two siRNA oligos, we observed a decrease in the expression of N-cadherin, ZEB1 and Slug. Coincident with the decrease in EMT-TFs, E-cadherin levels were elevated **(Fig. 5B)**. While there was no significant change in the *Slug* and *ZEB1* mRNA upon EPSTI1 knockdown **(Supplementary Fig. 2B)**, EPSTI1 disruption decreased N-cadherin mRNA expression over 50% in HCT116 and SW480 cells **(Fig. 5C)**. These results indicate that the switch of E-cadherin to N-cadherin expression promotes the progression of migratory and invasive behavior orchestrated by KSR1-EPSTI1 signaling in CRC cells.

**Figure 5.**
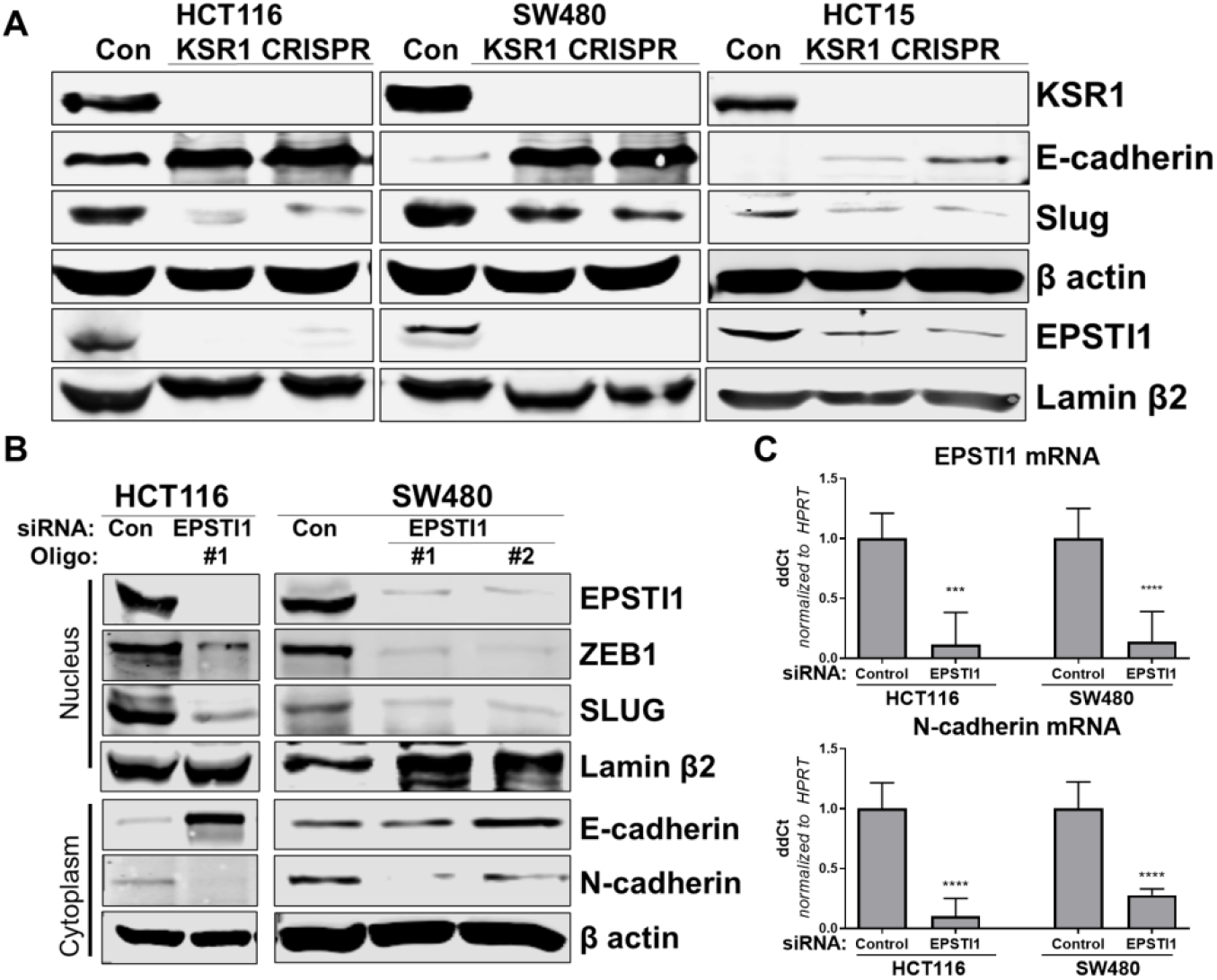
KSR1 and EPSTI1 promote cadherin switching. **(A)** Western blot analysis of the cell lysates prepared from control, and two clones of CRISPR-targeted HCT116, SW480, and HCT15 cells (KSR1 CRISPR) for the E-cadherin, Slug and EPSTI1. **(B)** Western blot of ZEB1, Slug, E-cadherin and N-cadherin in HCT116 and SW480 cells 72 hours following EPSTI1 knockdown. **(C)** RT-qPCR analysis of *EPSTI1* mRNA (upper) and *N-cadherin* (lower) following knockdown of EPSTI1 for 72 hours in HCT116 and SW480 cells. n=6; ***, P<0.001; ****, P<0.0001. Western blots shown in (A) and (B) and qPCR shown in (C) are representative of at least three independent experiments.

### EPSTI1 is necessary and sufficient for EMT in CRC cells

To determine the extent to which KSR1- and ERK-dependent EPSTI1 translation is critical to colon tumor cell growth and invasion, we expressed a MSCV-FLAG-EPSTI1-GFP construct in KSR1-CRISPR knockout HCT116, SW480, and HCT15 cells. CRISPR/Cas9-mediated deletion of KSR1 disrupted EPSTI1 expression, downregulated Slug and N-cadherin expression and elevated E-cadherin expression **(Fig. 6A)**. E-cadherin staining was absent in control CRC cells but evident at the cell membrane in KSR1 knockout cells **(Fig. 6B)**. Exogenous expression of EPSTI1 in cells lacking KSR1 restored the cadherin switch, by decreasing the expression of E-cadherin **(Fig. 6A and 6B)** and increasing N-cadherin levels comparable to control cells **(Fig. 6A)**. Suppression of E-cadherin and restoration of N-cadherin expression by the EPSTI1 transgene reestablished the ability of KSR1 knockout cells to migrate in monolayer culture **(Fig. 6C)** and invade through Matrigel®. Forced expression of EPSTI1 in these cells, increased the number of invading cells by over three-fold **(Fig. 6D)**. These data reveal that disabling the cadherin switch and inhibition of cell invasion by KSR1 disruption interrupts EPSTI1 translation, highlighting the pivotal role of this pathway for the induction of EMT in CRC cells.

**Figure 6.**
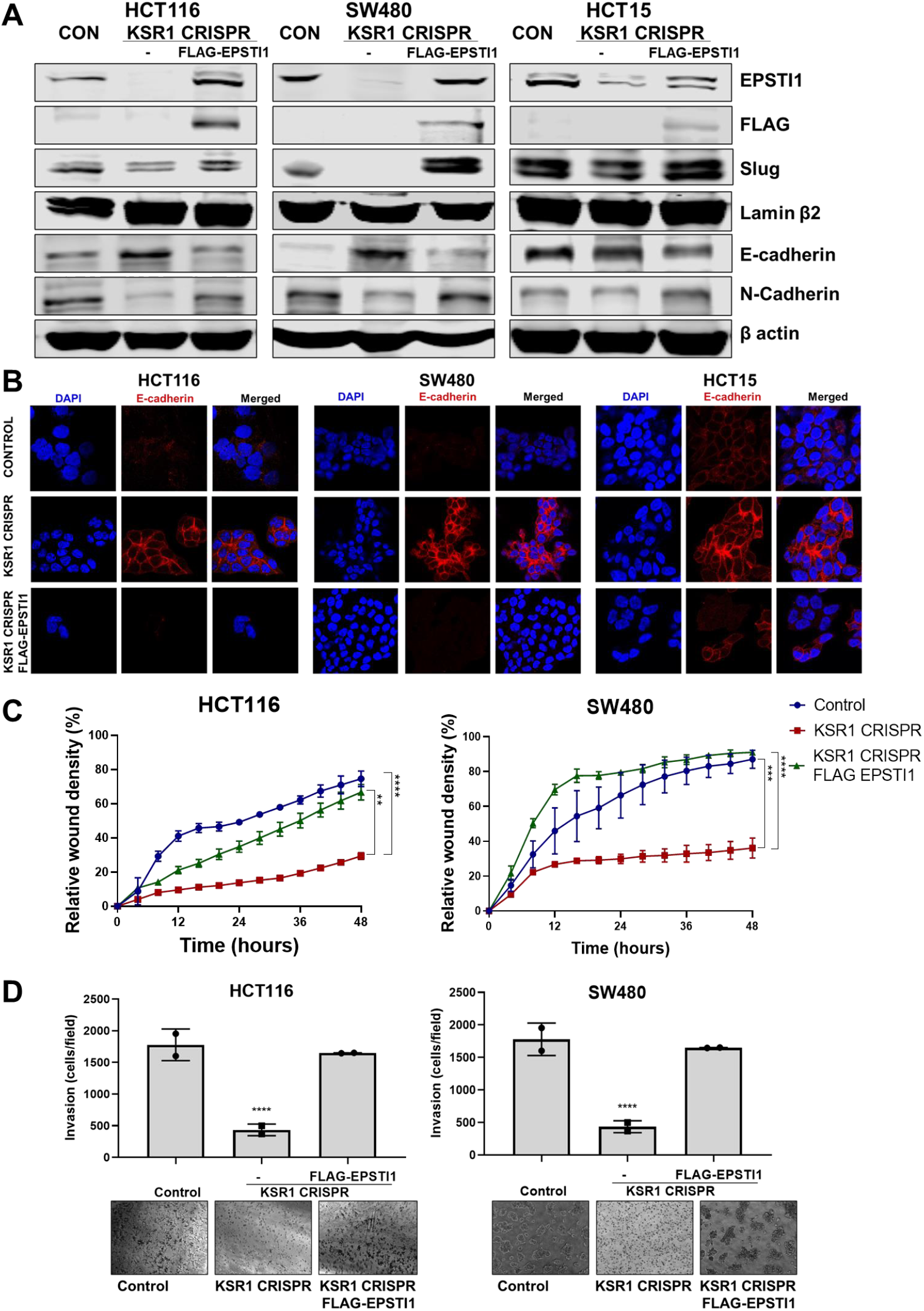
EPSTI1 rescues cadherin switching and invasive behavior to KSR1 knockout cells. **(A)** EPSTI1 protein expression was assessed by Western blotting in control, KSR1-targeted (KSR1 CRISPR) HCT116, SW480, and HCT15 cells with and without EPSTI1 (FLAG-EPSTI1) expression. Cells were lysed and probed for Slug, E-cadherin, N-cadherin, Lamin β2, and β actin. **(B)** Immunofluorescence staining for E-cadherin (Red) and DAPI (blue) in control or KSR1-targeted (KSR1 CRISPR) HCT116, SW480, and HCT15 cells with and without EPSTI1 (FLAG-EPSTI1) expression. **(C)** Control, CRIPSR-targeted (KSR1-CRISPR) and CRISPR-targeted HCT116 and SW480 cells expressing EPSTI1 (KSR1 CRISPR + FLAG-EPSTI1) were subjected to the 96-well IncuCyte scratch wound assay. The graph represents the time kinetics of percent wound density, calculated by IncuCyte ZOOM software, shown as mean ±SD, n=12; **, P < 0.005; ***, P < 0.001; ****, P<0.0001. Matched results were analyzed for statistical significance using one-way ANOVA with Dunnett’s posttest for multiple comparisons. **(D)** Control, CRISPR-targeted (KSR1 CRISPR) and CRISPR-targeted (E) HCT116 and (F) SW480 cells expressing EPSTI1 (KSR1 CRISPR + FLAG-EPSTI1) were subjected to Transwell migration assay through Matrigel^®^. The number of invaded cells per field were counted, (n=4); ****, P < 0.0001. Representative microscopic images of the respective cells following invasion through Matrigel® are shown.

### EPSTI1 re-expression reverses the KSR1-dependent growth inhibition and N-cadherin gene expression

Knockdown of EPSTI1 in HCT116 and SW480, decreased N-cadherin mRNA expression 50% **(Fig. 5C)**. Upon KSR1 depletion, N-cadherin mRNA decreased 32% in HCT116 and 89% in SW480 cells **(Fig. 7A)**. Ectopic expression of EPSTI1 in these cells restored the N-cadherin mRNA expression to levels observed in control SW480 cells, while in HCT116 KSR1 KO, forced EPSTI1 expression increased N-cadherin mRNA levels 3-fold above that seen in control HCT116 cells **(Fig. 7A)**. These data indicate that EPSTI1 mediates KSR1-dependent regulation of expression of N-cadherin mRNA to promote invasive behavior in colon cancer cells.

**Figure 7.**
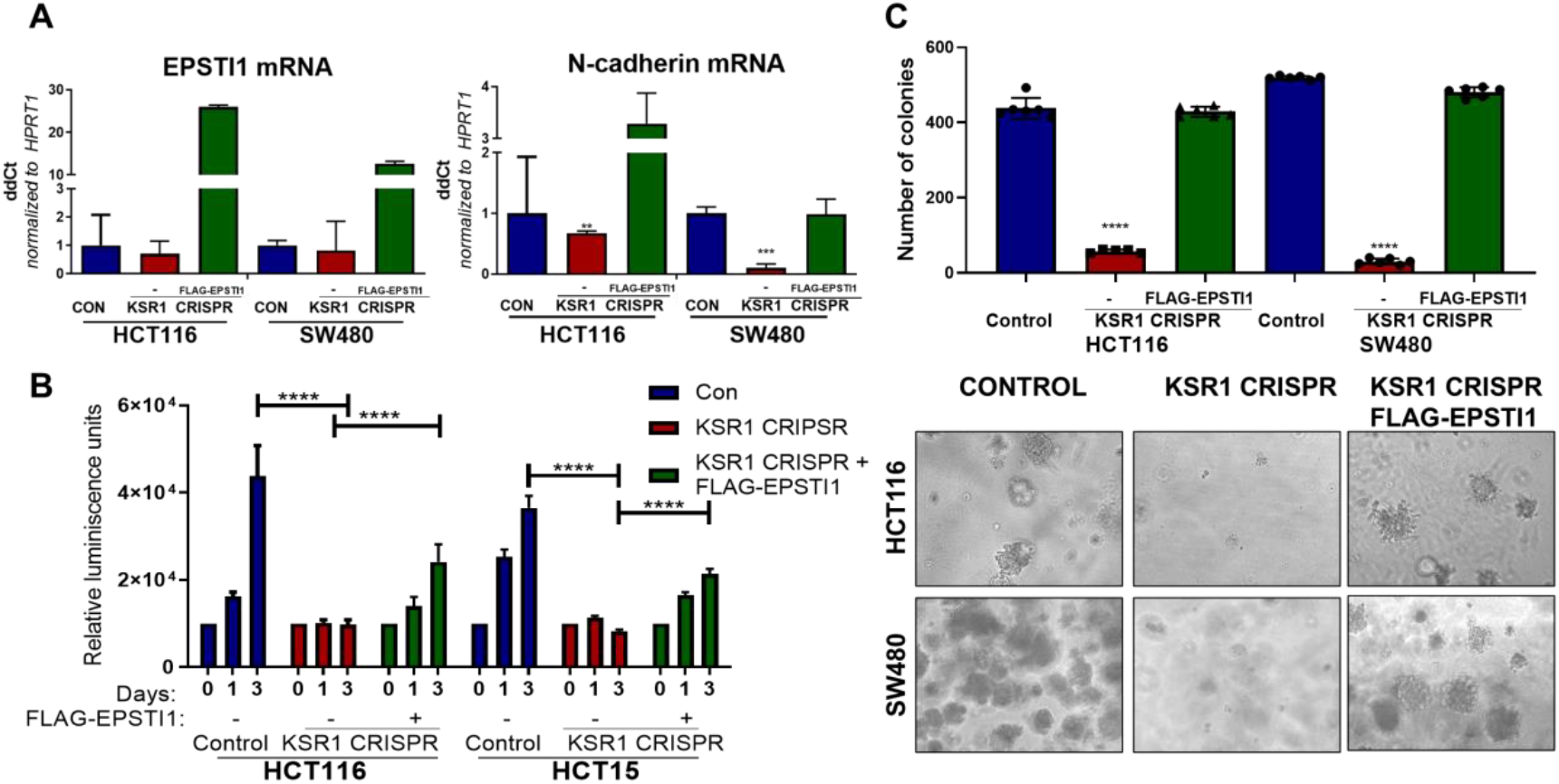
EPSTI1 expression in KSR1 KO cells induces N-cadherin mRNA expression and restores anchorage-independent growth. **(A)** RT-qPCR analysis of *EPSTI1* mRNA (left) and *N-cadherin* (right) in HCT116 and SW480 cells following KSR1 disruption with and without expression of EPSTI1 (FLAG-EPSTI1) in KSR1 KO cells. (n=3), **, P < 0.01***, P < 0.001 **(B)** KSR1 KO HCT116 and HCT15 cell viability (CellTiter-Glo) on poly-(HEMA)-coated plates at the indicated days with or without EPSTI1 (KSR1 CRISPR + EPSTI1) expression. The data are shown as relative luminescence units mean ± SD, (n=6); ****, P < 0.0001. The data were analyzed for statistical significance by one-way ANOVA followed by *t*-test. **(C)** Quantification of anchorage-independent colonies formed by KSR1 knockout HCT116 and SW480 cells with and without EPSTI1 expression (KSR1 CRISPR + FLAG-EPSTI1) after plating in soft agar. Representative photomicrographs of colonies from each cell line are shown. The data are illustrated as the number of colonies present after two weeks, (n=6) mean ± SD. ****, P < 0.0001. Data were analyzed for statistical significance one-way ANOVA followed by *t*-test.

The E- to N-cadherin switch promotes cancer cell survival following the loss of cell adhesion to the extracellular matrix (62, 63). KSR1 also promotes CRC cell survival when detached from a solid substrate (6, 17). To determine the extent to which EPSTI1 expression was sufficient to restore CRC cell viability in the absence of KSR1, we grew cells under anchorage-independent conditions either on Poly-(HEMA) **(Fig. 7B)** or on soft agar **(Fig. 7C)** following forced expression of EPSTI1 in HCT116, HCT15, and SW480 cells lacking KSR1. Anchorage-independent viability was measured over three days on poly-(HEMA) coated plates. Compared to control HCT116 and HCT15 cells, viability decreased approximately 75% in cells lacking KSR1. Ectopic expression of EPSTI1 restored viability to approximately 50% of control levels in both cell lines **(Fig. 7B)**. Similar to our previous findings (6, 55), KSR1 disruption hampered the ability of Ras transformed cells to form colonies on soft agar, the number of colonies formed in HCT116 and SW480 cells dramatically decreased by 75% in the absence of KSR1. Forced expression of EPSTI1 was sufficient to reverse the suppression of colony formation caused by KSR1 disruption to levels observed in control HCT116 and SW480 cells **(Fig. 7C)**. These results show that despite the absence of KSR1 to maintain and support cell growth, ectopic EPSTI1 expression was able to maintain anchorage-independent viability in CRC cells.

## Discussion

Persistent oncogenic reprogramming of transcription and translation during EMT grants migratory and invasive properties to tumor cells (22, 23). Multiple studies have established a relationship between oncogenic Ras-mediated ERK signaling and EMT, either through Ras or its downstream effector signaling pathways activating EMT-TFs (41, 43-47, 64). Silencing of Erbin, a tumor suppresser known to disrupt KSR1-RAF1 interaction, promoted cell migration and invasion of colon cancer cells, but did not identify the mechanism on how KSR1-dependent MAPK signaling affected EMT (65). Mediators of EMT activate cap-dependent translation initiation have been associated with increased aggressiveness and metastases of cancer cells, and we have shown that KSR1 can affect translation initiation (17, 48, 50, 66).

Our observations establish the novel role of the scaffold protein KSR1 promoting the preferential translation of an EMT-related gene, *EPSTI1*, and outline a mechanism for KSR1-dependent stimulation of EMT. Using gene-expression analysis of the polysome-bound mRNA, we discovered KSR1 and ERK increase the translational efficiency of *EPSTI1* mRNA. EPSTI1 mediates KSR1-dependent motility, invasion, and anchorage-independent growth coincident with its suppression of EMT-TF, Slug, elevating E-cadherin expression. EPSTI1 knockdown also decreased the expression of N-cadherin mRNA and protein. In the absence of KSR1, ectopic expression of EPSTI1 was sufficient to suppress E-cadherin expression, stimulate N-cadherin expression and enhance motility and invasive behavior. These data demonstrate that a KSR1- and ERK-regulated component is critical to the execution of the transcriptional program that drives interconversion between epithelial and mesenchymal phenotypes. These studies of post-transcriptional regulation and mRNA translation reveal the importance of expanding beyond gene expression analysis for detecting mechanisms underlying epithelial plasticity and tumorigenicity.

The association of EPSTI1 with tumor metastatic potential is supported by observations that *EPSTI1* is highly upregulated in invasive breast cancer tissues and suggested the role of EPSTI1 in promoting metastasis, tumorsphere formation, and stemness (51-53). Although the aberrant expression of EPSTI1 in breast cancer cells is well-established, there is little indication in the literature on the role of EPSTI1 to induce EMT, cancer invasion, and metastasis. The association of EPSTI1 induction of invasion in breast cancer cells was attributed to the increased expression of *Slug* and *Twist* mRNA and increased expression of fibronectin and α2β1 integrins (53). Another study suggested the interaction of EPSTI1 with valosin-containing protein (VCP) and the subsequent activation of NF-κB signaling contributed to the increased tumor invasion and metastasis (52). Future studies should evaluate the potential of EPSTI1 to directly affect N-cadherin and EMT-TF expression and assess the role of NF-κB signaling in EPSTI1-dependent CRC cell EMT.

Determining how KSR1- and ERK-dependent signaling promotes EPSTI1 translation should yield novel mechanisms underlying tumor cell metastatic behavior. We show that EPSTI1 mRNA is unchanged upon KSR1 disruption or ERK inhibition **(Figs. 1E and 2C)**, suggesting that KSR1 regulates EPSTI1 through post-transcriptional modifications enhancing its preferential loading onto the polysomes. Differential mRNA splicing is implicated in EMT-related processes and splicing regulatory factors have been implicated in the motility and invasive behavior of tumor cells (67, 68). One possibility is that KSR1 signaling promotes the splicing of EPSTI1 that promotes it’s the preferential translational contributing to increased motility and invasion.

Upon removal of KSR1 or EPSTI1, the tumor cells switch back from highly migratory and invasive EMT state to the epithelial state. However, the invasive property is not completely lost in KSR1/EPSTI1 disruption **(Fig. 4B)**, which could be attributed to other mesenchymal markers retained in the cells, such as vimentin **(Supplementary Fig. 2A)**. Investigating other EMT-related mRNAs that are preferentially translated in response to KSR1-scaffolded ERK signaling may reveal additional mRNAs that make previously unappreciated contributions to cell migration, invasion, and EMT. Constitutive KSR1 or EPSTI1 knockout yields developmentally normal mice (69-71). While KSR1 or EPSTI1 may not be essential to EMT during normal development, they may play a role in other EMT-dependent events such as wound healing where cells collectively migrate, differentiate, and re-epithelialize keratinocytes around and/or within the damaged site. If their role in EMT is exclusive to tumor cells it will reveal a key vulnerability for therapeutic evaluation. Further characterization of KSR1, EPSTI1 and the additional effectors repurposed by dysregulated translation in CRC should reveal additional novel mechanisms critical to CRC tumor survival and progression.

## Materials and Methods

### Cell culture

Colorectal cancer cell lines HCT116, HCT15 and SW480 were acquired from American Type Culture Collection (ATCC). The cells were cultured in Dulbecco’s modified Eagle’s medium (DMEM) containing high glucose with 10% fetal bovine serum (FBS) and grown at 37°C with ambient O_2_ and 5% CO_2_. Non-transformed immortalized human colon epithelial cell line (HCEC) was a gift from J. Shay (University of Texas [UT] Southwestern) and were grown and maintained as described previously (6, 72). HCECs were grown in a hypoxia chamber with 2% O_2_ and 5% CO_2_ at 37°C in 4 parts DMEM to 1 part medium 199 (Sigma-Aldrich #M4530) with 2% cosmic calf serum (GE Healthcare), 25 ng/mL EGF (R&D, Minneapolis, MN #236-EG), 1 µg/mL hydrocortisone (#H0888), 10 µg/mL insulin (#I550), 2 µg/mL transferrin (#T1428), 5 nM sodium selenite (Sigma-Aldrich #S5261), and 50 µg/mL gentamicin sulfate (Gibco #15750-060) as described previously (6). Normal and quadruple mutant AKPS (APC ^KO^/KRAS^G12D^/P53 ^KO^/SMAD4^KO^) tumor colon organoids obtained from the Living Organoid Biobank housed by Dr. Hans Clevers and cultured as described previously (60, 73). The normal organoids were cultured in medium containing advanced DMEM/F12 (Invitrogen #12634) with 50% WNT conditioned media (produced using stably transfected L cells), 20% R-spondin1, 10% Noggin, 1X B27 (Invitrogen #17504-044), 10 mM nicotinamide (Sigma-Aldrich #N0636), 1.25 mM N-acetylcysteine (Sigma-Aldrich #A9165-5G), 50 ng/mL EGF (Invitrogen #PMG8043), 5000 nM TGF-b type I receptor inhibitor A83-01 (Tocris #2939), 10 nM Prostaglandin E2 (Tocris #2296), 3 µM p38 inhibitor SB202190 (Sigma-Aldrich #S7067), and 100 µg/mL Primocin (Invivogen #ant-pm-1). The quadruple mutant AKPS organoids were grown in media lacking WNT conditioned media, R-spondin 1, noggin and EGF and containing 10 µM nutlin-3 (Sigma #675576-98-4).

### RNA interference

Approximately 500,000 cells were transfected using a final concentration of 20 nM EPSTI1 (J-015094-09-0020 and J-015094-12-0020) or non-targeting (D-001810-01-20 and D-001810-02-20) ON-TARGETplus siRNAs from GE Healthcare Dharmacon using 20 µL of Lipofectamine RNAiMAX (ThermoFisher #13778-150) and 500 µL OptiMEM (ThermoFisher #31985070). Cells were incubated for 72 hours before further analysis.

### Generation of KSR1 shRNA knockdown and KSR1 CRISPR/Cas9 knockout cell lines

A lentiviral pLKO.1-puro constructs targeting KSR1 and non-targeting control were transfected into HEK-293T cells using trans-lentiviral packaging system (ThermoFisher Scientific). The virus was collected, and the medium was replaced 48 hours post transfection. HCT116 and HCT15 cells were infected with virus with 8 µg/mL of Polybrene for several days. The population of cells with depleted KSR1 was selected with 10 µg/mL puromycin. The KSR1 knockdown was confirmed via Western Blotting.

pCAG-SpCas9-GFP-U6-gRNA was a gift from Jizhong Zou (Addgene plasmid #79144), KSR1 sgRNA and non-targeting control sgRNA was cloned into the pCas9 vector. Both the non-targeting control and sgKSR1 were transfected into HCT116, HCT15 and SW480 cells using PEI transfection as described previously (74). The GFP-positive cells were sorted 48-hours post transfection, and colonies were picked by placing sterile glass rings around individual colonies.

### Cell lysis and western blot analysis

Whole cell lysate was extracted in radioimmunoprecipitation assay (RIPA) buffer containing 50 mM Tris-HCl, 1% NP-40, 0.5% Na deoxycholate, 0.1% Na dodecyl sulfate, 150 mM NaCl, 2 mM EDTA, 2 mM EGTA, and 1X protease and phosphatase inhibitor cocktail (Halt, ThermoFisher Scientific #78440). Cytoplasmic and nuclear fractionation was performed using NE-PER™ Nuclear and Cytoplasmic Extraction Reagents (ThermoFisher Scientific #PI78835). The estimation of protein concentration was done using BCA protein assay (Promega #PI-23222, PI-23224). Samples were diluted using 1X sample buffer (4X stock, LI-COR #928-40004) with 100 mM dithiothreitol (DTT) (10X stock, 1mM, Sigma #D9779-5G). The protein was separated using 8-12 % SDS-PAGE and transferred to nitrocellulose membrane. The membrane was blocked with Odyssey TBS blocking buffer (LICOR-Biosciences #927-50003) for 45 minutes at room temperature, then incubated with primary antibodies *(Key Resources Table*) at least overnight at 4°C. IRDye 800CW and 680RD secondary antibodies (LI-COR Biosciences # 926-32211, # 926-68072) were diluted 1:10,000 in 0.1% TBS-Tween and imaged on the Odyssey Classic Scanner (LI-COR Biosciences).

### Polysome profiling

Cells were treated with 100 µg/mL cycloheximide (Sigma #C4859) on ice in PBS for 10 minutes. The cells were lysed with 10 mM HEPES, 100 mM KCL, 5 mM MgCl_2_, 100 µg/mL cycloheximide, 2 mM DTT, 1% Triton-X100, 2.5 µl RNaseOUT (ThermoFisher Scientific #10777019). The lysate were cleared by centrifugation for 10 minutes at 13,200rpm at 4°C. Approximately 200 µL of the total RNA was collected in a new RNAse-free microcentrifuge tube and the remaining supernatant was loaded onto a 15-45% sucrose gradient. The samples were spun at 37,500 rpm for 2 hours at 4°C in SW55Ti Beckman ultracentrifuge and separated on a gradient fractionation system to resolve the polysomes. Polysome profiles were identified at 260 nM using an absorbance detector. Gradient fractions were collected dropwise at 0.75mL/min. For RNAseq, the total RNA and RNA pooled from the polysome fraction (fractions 6-9) of three sets of independently isolated cells was isolated using RNAzol (Molecular Research Centre #RN 190) according to the manufacture’s protocol. RNA purity was evaluated by the UNMC DNA Sequencing Core using a BioAnalyzer.

### RNA-sequencing and analysis

RNA sequencing (RNAseq) was conducted by the UNMC DNA Sequencing Core. For RNA-seq, RNA was purified from three biological replicates of total and polysome-bound RNA from HCT116 and HCT15, control and KSR1 knockdown cells as previously described. Stranded RNA sequencing libraries were prepared as per manufactures’ protocol using TrueSeq mRNA protocol kit (Illumina) and 500 ng of the total RNA was used for each of the samples. Purified libraries were pooled at a 0.9 pM concentration and sequenced on an Illumina NextSeq550 instrument, using a 75 SR High-output flow cell, to obtain approximately 45 million single-end reads per sample. NGS short reads from RNA-seq experiments was downloaded from the HiSeq2500 server in FASTQ format. FastQC (http://www.bioinformatics.babraham.ac.uk/projects/fastqc/) was used to perform quality control checks on the *fastq* files that contain the raw short reads from sequencing. The reads were then mapped to the *Homo sapiens* (human) reference genome assembly GRCh38 (hg38) using STAR v2.7 alignment. The-*-quantMode GeneCounts* option in STAR 2.7 (75) was used to obtain the HTSeq counts per gene. Gencode v32 Gene Transfer Format (GTF) was used for the transcript/gene annotations. The output files were combined into a matrix using R. The gene counts were further used as input for downstream analysis using Anota2seq. The high-throughput sequencing data have been deposited in the Gene Expression Omnibus (GEO) database, www.ncbi.nlm.nih.gov/geo (accession no. GSE164492).

### Translational Efficiency

The altered levels of total mRNA can impact the changes in the pool of polysome-bound mRNA, leading to a spurious calculation translational efficiency (TE). Anota2seq (57) allows the quantification of actual changes in TE. TE was calculated using the R Bioconductor anota2Seq package for the HTSeq counts by first removing genes that did not contain expression values in more than 10% of the samples. 16,023 genes remained after this step. TMM normalization was further performed prior to log2 counts per million computation (CPM) using the voom function of the limma package using the anota2seqDataSetFromMatrix function (with parameters datatype?=?”RNAseq”, normalize?=?TRUE, transformation?=?”TMM-log2”). TE was calculated using the 2 × 2 factorial design model for the two cell lines (HCT116 and HCT15). Genes were considered significantly regulated at Adjusted p-value < 0.05 when passing filtering criteria (parameters for anota2seqSelSigGenes function) using Random variance Model [useRVM = TRUE], [selDeltaPT >log2(1.2)], [minSlopeTranslation >−1], [maxSlopeTranslation <2], [selDeltaTP >log2(1.2)], [minSlopeBuffering >−2] and [maxSlopeBuffering <1], [selDeltaP >log2(1)], [selDetaT >log2(1)]. The scatterplots were obtained using the anota2seqPlotFC function. The heatmaps were generated using the TE values for the two cell lines using the R Bioconductor ComplexHeatmap package.

### Anchorage-independent growth [poly-(HEMA)] assay

Poly-(HEMA) stock solution (10 mg/mL) was prepared by dissolving poly-(HEMA) (Sigma #3932-25G) in 95% ethanol at 37°C until fully dissolved (overnight). Ninety-six-well optical bottom plates (Thermo Scientific Nunc #165305) were coated in 200 µl of poly-(HEMA) solution and allowing it to evaporate. Cells were plated in complete growth medium of the poly-(HEMA) coated plates at a concentration of 10,000 cells/ 100 µL. Cell viability was measured at the indicated time points by the addition of CellTiter-Glo 2.0 reagent (Promega #G9242) and luminescence was measured (POLARstar Optima plate reader) according to the manufacturer’s protocol.

### Anchorage-independent growth (soft agar) assay

A total of 6000 cells were seeded in 1.6% NuSieve Agarose (Lonza #50081) to assess anchorage-independent growth according to the protocol of Fisher *et al*. (6). Colonies greater than 100 µm in diameter from 6 replicates per sample were counted, representative photomicrographs were taken after 10-14 days of incubation at 37°C and 5% CO2.

### RT-qPCR

Cells were harvested using 1 mL TRIzol (ThermoFisher Scientific #15596026) and RNA extraction was performed using RNeasy spin columns (Qiagen #74104). RNA was eluted with nuclease-free water. The RNA was quantified using a NanoDrop 2000 (Thermo Scientific) and Reverse Transcription (RT) was performed with 2 µg RNA per 40 µl reaction mixture using iScript Reverse Transcription Supermix (Bio-Rad #170-8891). RT-qPCR was performed using primers antibodies *(Key Resources Table*), and all targets were amplified using SsoAdvanced Universal SYBR green Supermix (Bio-Rad #1725271) with 40 cycles on a QuantStudio™ 3 (ThermoFisher Scientific). The analysis was performed using 2^-ΔΔC^_T_ method (76). For polysome gradients, the RNA levels were quantified from the cDNA using the standard curve method, summed across all fractions (3-8) and presented as a percentage of the total fractions.

### Cell migration (Scratch-test) assay

An *in vitro* scratch test were performed with the IncuCyte Zoom according to the manufacturer’s instructions. Approximately 35,000 cells were seeded onto a 96-well ImageLock plates (Essen BioScience #4379) and grown to 90-95% confluency. The scratches were created using WoundMaker (Essen BioScience #4563) in all the wells, after which the cells were washed with 1x PBS, and media without containing serum was replaced. Images of the cells were obtained every 20 minutes for a total duration of 72 hours using IncuCyte Kinetic Live Cell Imaging System (Essen BioScience) and analyzed using the IncuCyte Zoom software (Essen BioScience). Relative wound density was calculated as the percentage of spatial cell density inside the wound relative to the spatial density outside of the wound area at a given time point. The calculation of cell migration using this method, avoids false changes in cell density due to proliferation.

### Cell invasion (transwell) assay

Transwell inserts (24-well Millicell cell culture, #MCEP24H48) were coated with 50 µl of Matrigel® and allowed to solidify for 15-30 minutes. Approximately 20,000 stably generated knockout cells, or cells after 48 hours of transfection were plated in serum free media in the upper chamber of transwell insert. Cells were allowed to invade toward 10% serum containing media in the lower chamber for 24 hours, after which cells and gel in the upper chamber was gently removed with a sterile cotton applicator and the cells in the lower side of the insert was fixed with 3.7% formaldehyde for two minutes, permeabilized with 100% methanol for 20 minutes and stained with Giemsa for 15 minutes. The numbers of cells were counted using an inverted microscope at x20 magnification.

### Immunofluorescence assay

Cells were plated on glass coverslips to 70-80% confluence for 48 hours in growth media. Cells were fixed in 1% formaldehyde diluted in PBS for 15 minutes. The cells were rinsed three times with PBS for 5 minutes and coverslips were blocked for 1 hour with 1X PBS/ 5% goat serum/ 0.3% Triton™ X-100 and then incubated with E-cadherin antibody (#4A2) overnight. Cells were washed three times for 5 min with PBS and incubated in anti-mouse IgG Alexa Fluor® 555 Conjugate (Cell signaling #4409) at a dilution of 1:500 for 1 hour. Coverslips were rinsed three times for 5 min in PBS and briefly rinsed in distilled water prior to mounting in Prolong® Gold Antifade Reagent with DAPI (Cell signaling #8961). All Images were acquired using a Zeiss LSM-780 confocal microscope and processed using ZEISS ZEN 3.2 (blue edition) software.

### Key Resources Table

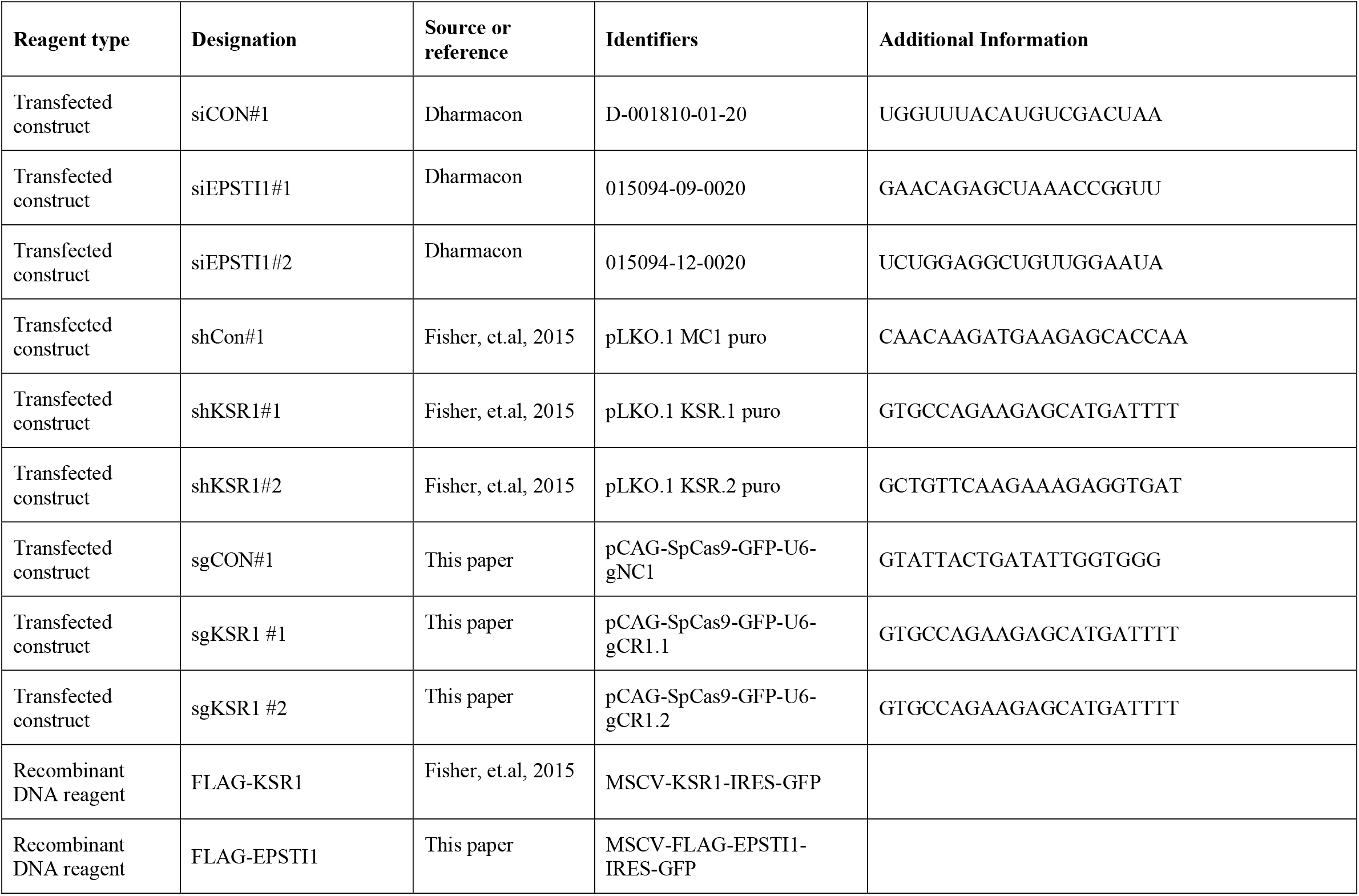
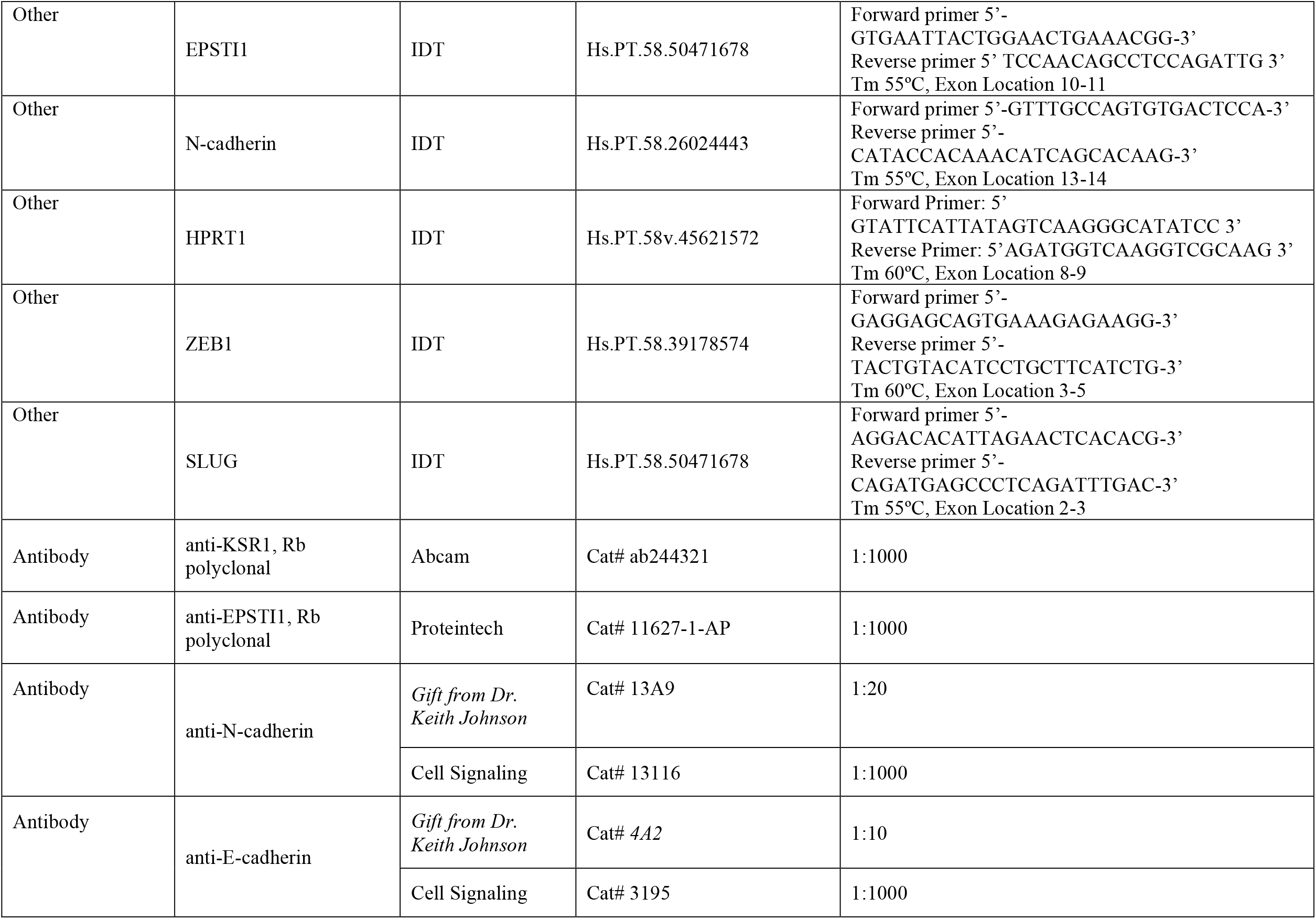
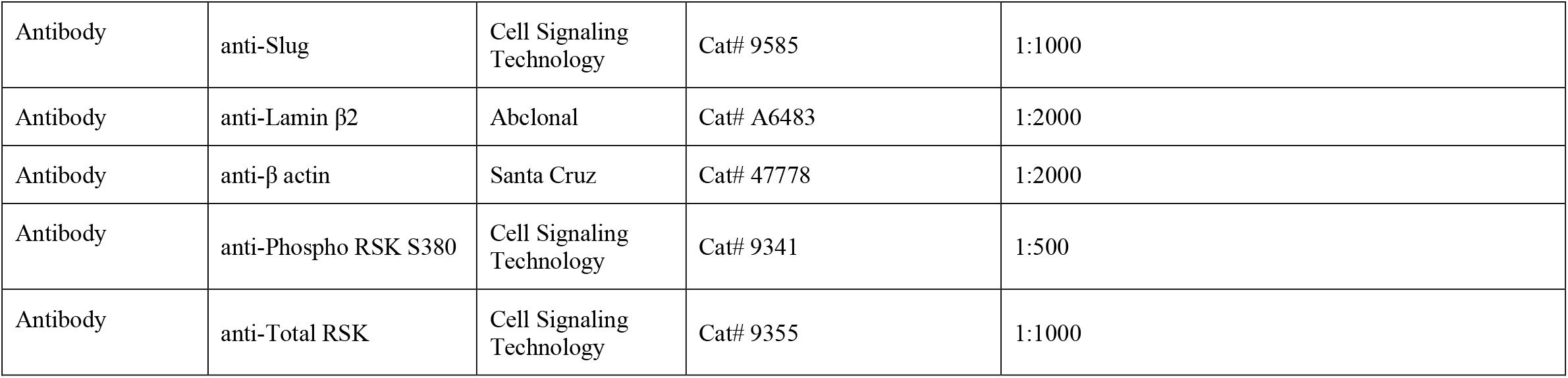

## Acknowledgements

We thank Dr. Xuan Zhang, and Dr. Kai Fu for their assistance with polysome profiling, Lisa E. Humphrey-Brattain for their assistance with the colon organoid culture, the UNMC Advanced Microscopy Core, and the UNMC Cell Analysis Facility. We declare no conflicts of interest.

---

**Supplementary Table 1:** Translational efficiency of mRNAs (58 decreased and 40 increased) upon KSR1 KD in HCT116 and HCT15 cells.

| No | Gene Symbol | translation.apvEff | apvRvmP |
| --- | --- | --- | --- |
| 1 | RN7SKP173 | -2.12 | 0.010526 |
| 2 | AC016074.2 | -1.93 | 0.022299 |
| 3 | FKBP2 | -1.63 | 0.000661 |
| 4 | RP11-89K21.1 | -1.61 | 0.013317 |
| 5 | GLT8D2 | -1.53 | 0.025778 |
| 6 | AC034139.1 | -1.41 | 0.041679 |
| 7 | EPSTI1 | -1.40 | 0.003656 |
| 8 | RUNX3 | -1.39 | 0.024100 |
| 9 | RP11-462G12.1 | -1.38 | 0.008471 |
| 10 | KIAA1377 | -1.26 | 0.025489 |
| 11 | AC074289.1 | -1.26 | 0.009760 |
| 12 | MURC | -1.25 | 0.006237 |
| 13 | RP11-119F7.5 | -1.24 | 0.014357 |
| 14 | WNT9A | -1.23 | 0.025868 |
| 15 | RP11-417L19.4 | -1.21 | 0.018054 |
| 16 | GRM2 | -1.19 | 0.020653 |
| 17 | DLL1 | -1.18 | 0.010732 |
| 18 | CTD-2619J13.17 | -1.12 | 0.004376 |
| 19 | RNF112 | -1.03 | 0.011236 |
| 20 | HOXA10-AS | -1.02 | 0.006914 |
| 21 | HOXC8 | -1.02 | 0.023323 |
| 22 | AC245140.3 | -1.01 | 0.031525 |
| 23 | FRMPD1 | -0.99 | 0.020411 |
| 24 | LRRN4 | -0.95 | 0.011565 |
| 25 | COL13A1 | -0.93 | 0.005897 |
| 26 | CTC-453G23.8 | -0.93 | 0.004409 |
| 27 | PTPRN2 | -0.92 | 0.002917 |
| 28 | BEND3P3 | -0.91 | 0.010644 |
| 29 | RP11-448G15.3 | -0.91 | 0.022294 |
| 30 | MT-TF | -0.90 | 0.022067 |
| 31 | MRC2 | -0.88 | 0.025039 |
| 32 | P2RY1 | -0.87 | 0.019079 |
| 33 | MGAT2 | -0.87 | 0.022811 |
| 34 | LHX9 | -0.86 | 0.012179 |
| 35 | MTMR9 | -0.82 | 0.030426 |
| 36 | USP2 | -0.81 | 0.011305 |
| 37 | IFIT3 | -0.81 | 0.021296 |
| 38 | RP11-218C14.8 | -0.79 | 0.005042 |
| 39 | ABCG2 | -0.76 | 0.009184 |
| 40 | EGR2 | -0.76 | 0.015341 |
| 41 | RP11-713C19.2 | -0.76 | 0.033777 |
| 42 | ZFP2 | -0.75 | 0.008896 |
| 43 | RASD1 | -0.73 | 0.024515 |
| 44 | RP11-757G1.6 | -0.70 | 0.031518 |
| 45 | KIAA0825 | -0.70 | 0.014460 |
| 46 | SOBP | -0.69 | 0.035367 |
| 47 | CEP19 | -0.67 | 0.013100 |
| 48 | AQP1 | -0.67 | 0.013263 |
| 49 | TUBBP1 | -0.64 | 0.011810 |
| 50 | SEPTIN3 | -0.62 | 0.038409 |
| 51 | RP5-968P14.2 | -0.58 | 0.041242 |
| 52 | RP5-1139B12.3 | -0.57 | 0.018554 |
| 53 | HAL | -0.56 | 0.038409 |
| 54 | VANGL2 | -0.55 | 0.026444 |
| 55 | FEZF1 | -0.54 | 0.024427 |
| 56 | AC009802.1 | -0.53 | 0.003870 |
| 57 | GALNT12 | -0.53 | 0.027702 |
| 58 | C3orf80 | -0.52 | 0.024221 |
| 59 | AUH | 0.51 | 0.049181 |
| 60 | SLC11A1 | 0.51 | 0.033908 |
| 61 | RBM8B | 0.54 | 0.041679 |
| 62 | MEF2BNB | 0.54 | 0.042614 |
| 63 | APOBEC3H | 0.54 | 0.027176 |
| 64 | BBC3 | 0.56 | 0.006004 |
| 65 | HIST1H2AC | 0.59 | 0.024407 |
| 66 | MRPS6 | 0.60 | 0.022841 |

| No | Gene_Symbol | translation.apvEff | apvRvmP |
| --- | --- | --- | --- |
| 67 | ARHGAP30 | 0.62 | 0.032923 |
| 68 | ACTRT3 | 0.67 | 0.008309 |
| 69 | VAMP5 | 0.69 | 0.023729 |
| 70 | RP11-432J22.2 | 0.69 | 0.023571 |
| 71 | EFCAB6 | 0.73 | 0.005135 |
| 72 | SUMO2P17 | 0.74 | 0.003038 |
| 73 | PRAC2 | 0.76 | 0.025470 |
| 74 | SH2D6 | 0.77 | 0.033212 |
| 75 | AC073072.7 | 0.80 | 0.021478 |
| 76 | AC005932.1 | 0.81 | 0.031525 |
| 77 | RP11-54O7.18 | 0.82 | 0.015558 |
| 78 | CTC-490E21.10 | 0.84 | 0.022299 |
| 79 | AC005281.1 | 0.86 | 0.026496 |
| 80 | RP11-355B11.2 | 0.90 | 0.005146 |
| 81 | NFATC2 | 0.93 | 0.022930 |
| 82 | AC008155.1 | 0.94 | 0.019878 |
| 83 | HSD52 | 1.00 | 0.006602 |
| 84 | RP11-486I11.2 | 1.02 | 0.008414 |
| 85 | RP11-6B6.3 | 1.05 | 0.002162 |
| 86 | ZNF233 | 1.05 | 0.018509 |
| 87 | AP002813.1 | 1.06 | 0.008471 |
| 88 | EIF4A1P7 | 1.07 | 0.003762 |
| 89 | RP11-503N18.1 | 1.09 | 0.020907 |
| 90 | COX7C | 1.09 | 0.001558 |
| 91 | CTD-2342J14.6 | 1.10 | 0.002788 |
| 92 | TCP10L | 1.10 | 0.007539 |
| 93 | RP11-353N14.4 | 1.23 | 0.003870 |
| 94 | RP1-140A9.1 | 1.28 | 0.015622 |
| 95 | COX7CP1 | 1.32 | 0.001676 |
| 96 | RP11-663P9.2 | 1.33 | 0.004750 |
| 97 | RP11-85A1.3 | 1.37 | 0.005673 |
| 98 | SLC51B | 1.46 | 0.023832 |

**Supplementary Table 2:**
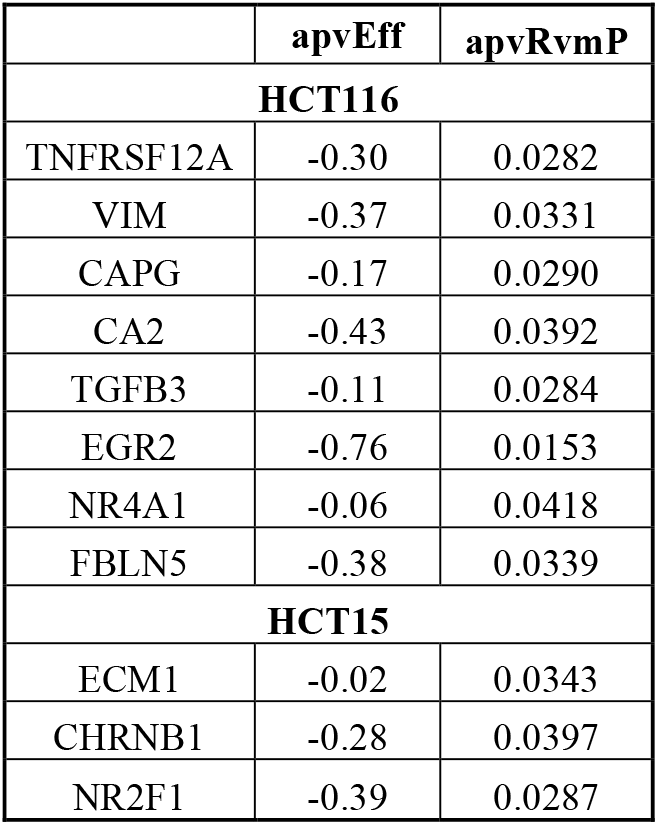
mRNAs translated in KSR1-dependent manner predicted in mesenchymal-up signature identified by GSEA.

**Figure S1.**
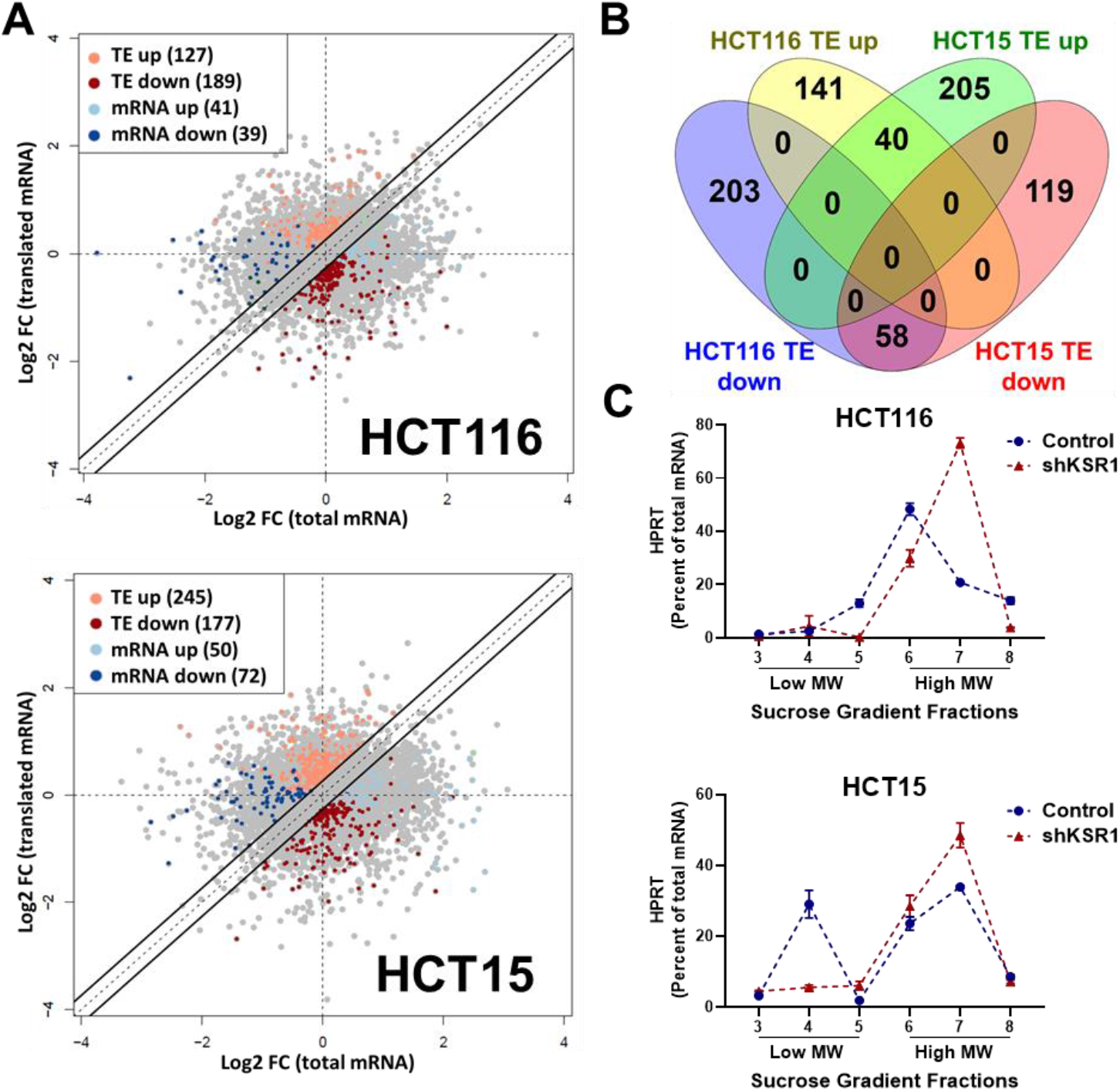
EPSTI1 is translationally regulated by KSR1. **(A)** Scatter plot of polysome-associated mRNA to total mRNA log2 fold-changes upon KSR1 knockdown in HCT116 (top) and HCT15 (bottom) with RNA-seq. The statistically significant genes in the absence of KSR1 are classified into four groups with a fold change (|log_2_FC| > 1.2) and p-value < 0.05. The number of mRNAs with a change in TE (orange and red) are indicated (n=3 for each cell line). TE, translational efficiency **(B)** Differential gene expression analysis comparing genes whose TE is changed upon KSR1-knockdown in HCT116 and HCT15 **(C)** RT-qPCR analysis of *HPRT* mRNA levels isolated from sucrose gradient fractions of the control and KSR1 knockdown HCT116 and HCT15 cells. Fractions 3-5 (low MW) and 6-8 (high MW) are plotted for the control and KSR1 knockdown state with values corresponding to the percentage of total mRNA across these fractions. Experiments shown in (A-C) are representative of three independent experiments.

**Figure S2.**
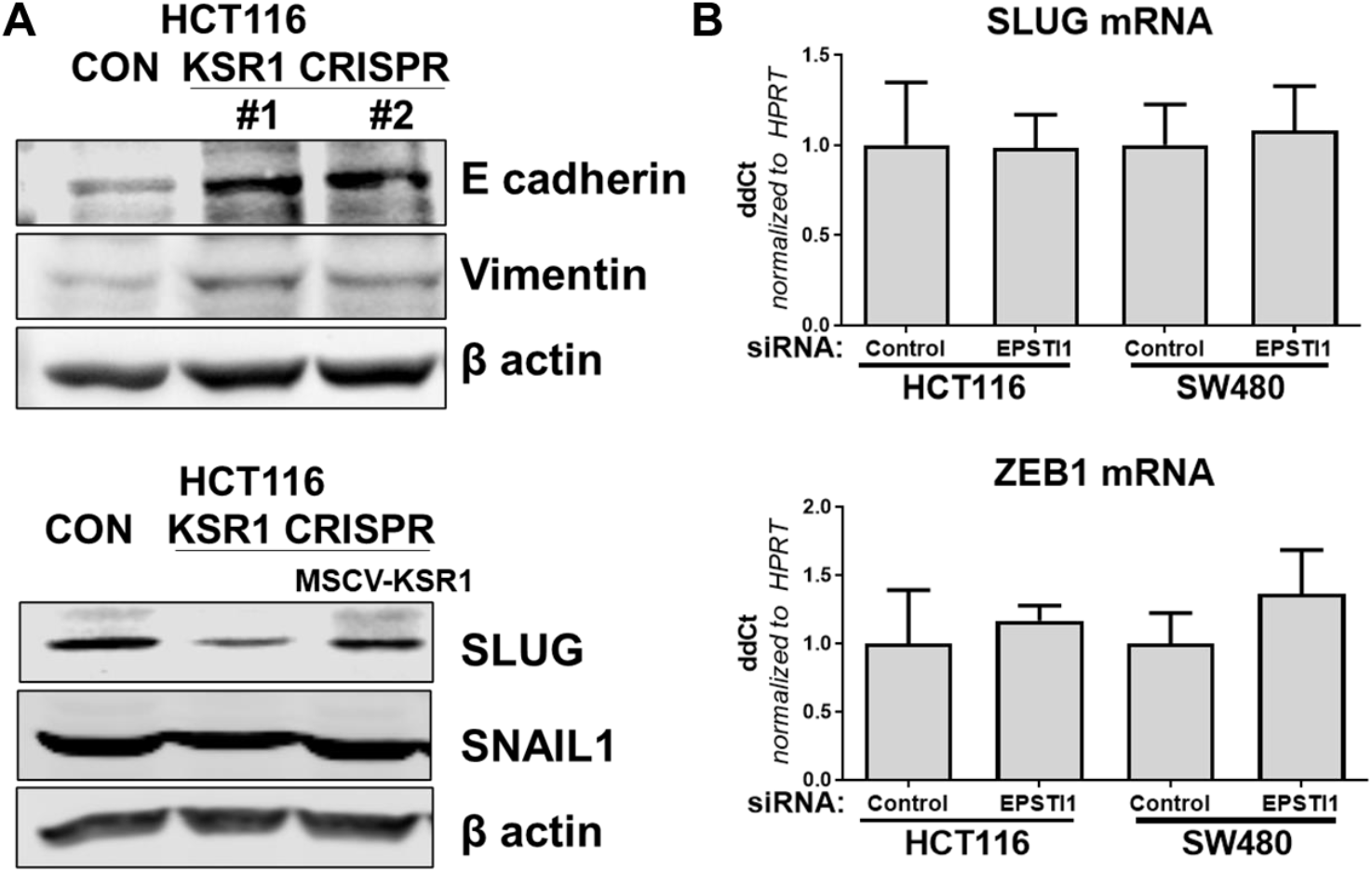
KSR1 and EPSTI1 promote the cadherin switch. **(A)** Western blot analysis of the cell lysates prepared from control, and two clones of CRISPR-targeted HCT116 (KSR1 CRISPR) for (Top-upper) E-cadherin and Vimentin, and (bottom-lower) Cell lysates prepared from control, CRISPR-targeted (KSR1 CRISPR) and CRISPR-targeted HCT116 cells expressing KSR1 (MSCV-KSR1) analyzed for Slug and Snail. **(B)** RT-qPCR analysis of Slug mRNA (top-upper) and ZEB1 (bottom-lower) following knockdown of EPSTI1 for 72 hours in HCT116 and SW480 cells.

## References

1. Morrison DK, Davis RJ. Regulation of MAP kinase signaling modules by scaffold proteins in mammals. Annu Rev Cell Dev Biol. 2003;19:91–118.

2. Pawson T, Scott JD. Signaling through scaffold, anchoring, and adaptor proteins. Science. 1997;278(5346):2075–80.

3. Kortum RL, Lewis RE. The Molecular Scaffold KSR1 Regulates the Proliferative and Oncogenic Potential of Cells. Molecular and Cellular Biology. 2004;24(10):4407–16.

4. Nguyen A, Burack WR, Stock JL, Kortum R, Chaika OV, Afkarian M, et al. Kinase suppressor of Ras (KSR) is a scaffold which facilitates mitogen-activated protein kinase activation in vivo. Molecular and Cellular Biology. 2002;22(9):3035–45.

5. Fisher KW, Das B, Kortum RL, Chaika OV, Lewis RE. Kinase suppressor of ras 1 (KSR1) regulates PGC1α and estrogen-related receptor α to promote oncogenic Ras-dependent anchorage-independent growth. Molecular and Cellular Biology. 2011;31(12):2453–61.

6. Fisher KW, Das B, Kim HS, Clymer BK, Gehring D, Smith DR, et al. AMPK Promotes Aberrant PGC1β Expression To Support Human Colon Tumor Cell Survival. Molecular and Cellular Biology. 2015;35(22):3866–79.

7. Morrison MMM, Daniel AR, Deborah K. Signaling dynamics of the KSR1 scaffold complex. 2009.

8. Haigis KM. KRAS Alleles: The Devil Is in the Detail. Trends Cancer. 2017;3(10):686–97.

9. Serebriiskii IG, Connelly C, Frampton G, Newberg J, Cooke M, Miller V, et al. Comprehensive characterization of RAS mutations in colon and rectal cancers in old and young patients. Nat Commun. 2019;10(1):3722.

10. Schmitz KJ, Wohlschlaeger J, Alakus H, Bohr J, Stauder MA, Worm K, et al. Activation of extracellular regulated kinases (ERK1/2) but not AKT predicts poor prognosis in colorectal carcinoma and is associated with k-ras mutations. Virchows Arch. 2007;450(2):151–9.

11. Brandt R, Sell T, Luthen M, Uhlitz F, Klinger B, Riemer P, et al. Cell type-dependent differential activation of ERK by oncogenic KRAS in colon cancer and intestinal epithelium. Nat Commun. 2019;10(1):2919.

12. Chu J, Cargnello M, Topisirovic I, Pelletier J. Translation Initiation Factors: Reprogramming Protein Synthesis in Cancer. Trends Cell Biol. 2016;26(12):918–33.

13. Avdulov S, Li S, Michalek V, Burrichter D, Peterson M, Perlman DM, et al. Activation of translation complex eIF4F is essential for the genesis and maintenance of the malignant phenotype in human mammary epithelial cells. Cancer Cell. 2004;5(6):553–63.

14. Truitt Morgan L, Conn Crystal S, Shi Z, Pang X, Tokuyasu T, Coady Alison M, et al. Differential Requirements for eIF4E Dose in Normal Development and Cancer. Cell. 2015;162(1):59–71.

15. Pelletier J, Graff J, Ruggero D, Sonenberg N. Targeting the eIF4F translation initiation complex: a critical nexus for cancer development. Cancer Research. 2015;75(2):250–63.

16. Truitt ML, Ruggero D. New frontiers in translational control of the cancer genome. Nat Rev Cancer. 2016;16(5):288–304.

17. McCall JL, Gehring D, Clymer BK, Fisher KW, Das B, Kelly DL, et al. KSR1 and EPHB4 Regulate Myc and PGC1β To Promote Survival of Human Colon Tumors. Molecular and Cellular Biology. 2016;36(17):2246–61.

18. Neilsen BK, Frodyma DE, McCall JL, Fisher KW, Lewis RE. ERK-mediated TIMELESS expression suppresses G2/M arrest in colon cancer cells. PLOS ONE. 2019;14(1):e0209224.

19. Ye X, Weinberg RA. Epithelial-Mesenchymal Plasticity: A Central Regulator of Cancer Progression. Trends Cell Biol. 2015;25(11):675–86.

20. Nieto MA. Epithelial plasticity: a common theme in embryonic and cancer cells. Science. 2013;342(6159):1234850.

21. Thiery JP, Acloque H, Huang RY, Nieto MA. Epithelial-mesenchymal transitions in development and disease. Cell. 2009;139(5):871–90.

22. Dongre A, Weinberg RA. New insights into the mechanisms of epithelial-mesenchymal transition and implications for cancer. Nat Rev Mol Cell Biol. 2019;20(2):69–84.

23. Nieto MA, Huang RY, Jackson RA, Thiery JP. EMT: 2016. Cell. 2016;166(1):21–45.

24. Thiery JP. Epithelial-mesenchymal transitions in tumour progression. Nat Rev Cancer. 2002;2(6):442–54.

25. Oda T, Kanai Y, Oyama T, Yoshiura K, Shimoyama Y, Birchmeier W, et al. Ecadherin gene mutations in human gastric carcinoma cell lines. Proc Natl Acad Sci U S A. 1994;91(5):1858–62.

26. Frixen UH, Behrens J, Sachs M, Eberle G, Voss B, Warda A, et al. E-cadherin-mediated cell-cell adhesion prevents invasiveness of human carcinoma cells. J Cell Biol. 1991;113(1):173–85.

27. Jolly MK, Ware KE, Gilja S, Somarelli JA, Levine H. EMT and MET: necessary or permissive for metastasis? Mol Oncol. 2017;11(7):755–69.

28. Nieman MT, Prudoff RS, Johnson KR, Wheelock MJ. N-cadherin promotes motility in human breast cancer cells regardless of their E-cadherin expression. J Cell Biol. 1999;147(3):631–44.

29. Liu CC, Cai DL, Sun F, Wu ZH, Yue B, Zhao SL, et al. FERMT1 mediates epithelial-mesenchymal transition to promote colon cancer metastasis via modulation of beta-catenin transcriptional activity. Oncogene. 2017;36(13):1779–92.

30. Suyama K, Shapiro I, Guttman M, Hazan RB. A signaling pathway leading to metastasis is controlled by N-cadherin and the FGF receptor. Cancer Cell. 2002;2(4):301–14.

31. Rosivatz E, Becker I, Bamba M, Schott C, Diebold J, Mayr D, et al. Neoexpression of N-cadherin in E-cadherin positive colon cancers. Int J Cancer. 2004;111(5):711–9.

32. Okubo K, Uenosono Y, Arigami T, Yanagita S, Matsushita D, Kijima T, et al. Clinical significance of altering epithelial-mesenchymal transition in metastatic lymph nodes of gastric cancer. Gastric Cancer. 2017;20(5):802–10.

33. Sadot E, Simcha I, Shtutman M, Ben-Ze’ev A, Geiger B. Inhibition of beta-catenin-mediated transactivation by cadherin derivatives. Proc Natl Acad Sci U S A. 1998;95(26):15339–44.

34. Loh CY, Chai JY, Tang TF, Wong WF, Sethi G, Shanmugam MK, et al. The E-Cadherin and N-Cadherin Switch in Epithelial-to-Mesenchymal Transition: Signaling, Therapeutic Implications, and Challenges. Cells. 2019;8(10).

35. Scarpa E, Szabo A, Bibonne A, Theveneau E, Parsons M, Mayor R. Cadherin Switch during EMT in Neural Crest Cells Leads to Contact Inhibition of Locomotion via Repolarization of Forces. Dev Cell. 2015;34(4):421–34.

36. Hulit J, Suyama K, Chung S, Keren R, Agiostratidou G, Shan W, et al. N-cadherin signaling potentiates mammary tumor metastasis via enhanced extracellular signal-regulated kinase activation. Cancer Res. 2007;67(7):3106–16.

37. Wheelock MJ, Shintani Y, Maeda M, Fukumoto Y, Johnson KR. Cadherin switching. J Cell Sci. 2008;121(Pt 6):727–35.

38. Tomita K, van Bokhoven A, van Leenders GJ, Ruijter ET, Jansen CF, Bussemakers MJ, et al. Cadherin switching in human prostate cancer progression. Cancer Res. 2000;60(13):3650–4.

39. Maeda M, Johnson KR, Wheelock MJ. Cadherin switching: essential for behavioral but not morphological changes during an epithelium-to-mesenchyme transition. J Cell Sci. 2005;118(Pt 5):873–87.

40. Araki K, Shimura T, Suzuki H, Tsutsumi S, Wada W, Yajima T, et al. E/N-cadherin switch mediates cancer progression via TGF-beta-induced epithelial-to-mesenchymal transition in extrahepatic cholangiocarcinoma. Br J Cancer. 2011;105(12):1885–93.

41. Shin S, Dimitri CA, Yoon S-O, Dowdle W, Blenis J. ERK2 but not ERK1 induces epithelial-to-mesenchymal transformation via DEF motif-dependent signaling events. Molecular Cell. 2010;38(1):114–27.

42. Shin S, Buel GR, Nagiec MJ, Han MJ, Roux PP, Blenis J, et al. ERK2 regulates epithelial-to-mesenchymal plasticity through DOCK10-dependent Rac1/FoxO1 activation. Proc Natl Acad Sci U S A. 2019;116(8):2967–76.

43. Andreolas C, Kalogeropoulou M, Voulgari A, Pintzas A. Fra-1 regulates vimentin during Ha-RAS-induced epithelial mesenchymal transition in human colon carcinoma cells. Int J Cancer. 2008;122(8):1745–56.

44. Liu Y, Sanchez-Tillo E, Lu X, Huang L, Clem B, Telang S, et al. The ZEB1 transcription factor acts in a negative feedback loop with miR200 downstream of Ras and Rb1 to regulate Bmi1 expression. J Biol Chem. 2014;289(7):4116–25.

45. Wong CE, Yu JS, Quigley DA, To MD, Jen KY, Huang PY, et al. Inflammation and Hras signaling control epithelial-mesenchymal transition during skin tumor progression. Genes Dev. 2013;27(6):670–82.

46. Wang Y, Ngo VN, Marani M, Yang Y, Wright G, Staudt LM, et al. Critical role for transcriptional repressor Snail2 in transformation by oncogenic RAS in colorectal carcinoma cells. Oncogene. 2010;29(33):4658–70.

47. Lemieux E, Bergeron S, Durand V, Asselin C, Saucier C, Rivard N. Constitutively active MEK1 is sufficient to induce epithelial-to-mesenchymal transition in intestinal epithelial cells and to promote tumor invasion and metastasis. Int J Cancer. 2009;125(7):1575–86.

48. Jechlinger M, Grunert S, Tamir IH, Janda E, Ludemann S, Waerner T, et al. Expression profiling of epithelial plasticity in tumor progression. Oncogene. 2003;22(46):7155–69.

49. Aiello NM, Maddipati R, Norgard RJ, Balli D, Li J, Yuan S, et al. EMT Subtype Influences Epithelial Plasticity and Mode of Cell Migration. Dev Cell. 2018;45(6):681–95 e4.

50. Waerner T, Alacakaptan M, Tamir I, Oberauer R, Gal A, Brabletz T, et al. ILEI: a cytokine essential for EMT, tumor formation, and late events in metastasis in epithelial cells. Cancer Cell. 2006;10(3):227–39.

51. Nielsen HL, Ronnov-Jessen L, Villadsen R, Petersen OW. Identification of EPSTI1, a novel gene induced by epithelial-stromal interaction in human breast cancer. Genomics. 2002;79(5):703–10.

52. Li T, Lu H, Shen C, Lahiri SK, Wason MS, Mukherjee D, et al. Identification of epithelial stromal interaction 1 as a novel effector downstream of Krüppel-like factor 8 in breast cancer invasion and metastasis. Oncogene. 2014;33(39):4746–55.

53. de Neergaard M, Kim J, Villadsen R, Fridriksdottir AJ, Rank F, Timmermans-Wielenga V, et al. Epithelial-Stromal Interaction 1 (EPSTI1) Substitutes for Peritumoral Fibroblasts in the Tumor Microenvironment. Am J Pathol. 1762010. p. 1229–40.

54. Roux PP, Shahbazian D, Vu H, Holz MK, Cohen MS, Taunton J, et al. RAS/ERK signaling promotes site-specific ribosomal protein S6 phosphorylation via RSK and stimulates cap-dependent translation. J Biol Chem. 2007;282(19):14056–64.

55. Kortum RL, Johnson HJ, Costanzo DL, Volle DJ, Razidlo GL, Fusello AM, et al. The molecular scaffold kinase suppressor of Ras 1 is a modifier of RasV12-induced and replicative senescence. Molecular and Cellular Biology. 2006;26(6):2202–14.

56. King HA, Gerber AP. Translatome profiling: methods for genome-scale analysis of mRNA translation. Brief Funct Genomics. 2016;15(1):22–31.

57. Oertlin C, Lorent J, Murie C, Furic L, Topisirovic I, Larsson O. Generally applicable transcriptome-wide analysis of translation using anota2seq. Nucleic Acids Res. 2019;47(12):e70.

58. Subramanian A, Tamayo P, Mootha VK, Mukherjee S, Ebert BL, Gillette MA, et al. Gene set enrichment analysis: A knowledge-based approach for interpreting genome-wide expression profiles. Proceedings of the National Academy of Sciences. 2005;102(43):15545–50.

59. Morris EJ, Jha S, Restaino CR, Dayananth P, Zhu H, Cooper A, et al. Discovery of a novel ERK inhibitor with activity in models of acquired resistance to BRAF and MEK inhibitors. Cancer Discovery. 2013;3(7):742–50.

60. Drost J, van Jaarsveld RH, Ponsioen B, Zimberlin C, van Boxtel R, Buijs A, et al. Sequential cancer mutations in cultured human intestinal stem cells. Nature. 2015;521(7550):43–7.

61. Johnston ST, Shah ET, Chopin LK, Sean McElwain DL, Simpson MJ. Estimating cell diffusivity and cell proliferation rate by interpreting IncuCyte ZOOM assay data using the Fisher-Kolmogorov model. BMC Syst Biol. 2015;9:38.

62. Derksen PW, Liu X, Saridin F, van der Gulden H, Zevenhoven J, Evers B, et al. Somatic inactivation of E-cadherin and p53 in mice leads to metastatic lobular mammary carcinoma through induction of anoikis resistance and angiogenesis. Cancer Cell. 2006;10(5):437–49.

63. Onder TT, Gupta PB, Mani SA, Yang J, Lander ES, Weinberg RA. Loss of E-cadherin promotes metastasis via multiple downstream transcriptional pathways. Cancer Res. 2008;68(10):3645–54.

64. Diesch J, Sanij E, Gilan O, Love C, Tran H, Fleming NI, et al. Widespread FRA1-dependent control of mesenchymal transdifferentiation programs in colorectal cancer cells. PLoS One. 2014;9(3):e88950.

65. Stevens PD, Wen Y-A, Xiong X, Zaytseva YY, Li AT, Wang C, et al. Erbin Suppresses KSR1-Mediated RAS/RAF Signaling and Tumorigenesis in Colorectal Cancer. Cancer Research. 2018;78(17):4839–52.

66. Prakash V, Carson BB, Feenstra JM, Dass RA, Sekyrova P, Hoshino A, et al. Ribosome biogenesis during cell cycle arrest fuels EMT in development and disease. Nat Commun. 2019;10(1):2110.

67. Park S, Brugiolo M, Akerman M, Das S, Urbanski L, Geier A, et al. Differential Functions of Splicing Factors in Mammary Transformation and Breast Cancer Metastasis. Cell Rep. 2019;29(9):2672–88 e7.

68. Pradella D, Naro C, Sette C, Ghigna C. EMT and stemness: flexible processes tuned by alternative splicing in development and cancer progression. Mol Cancer. 2017;16(1):8.

69. Lozano J, Xing R, Cai Z, Jensen HL, Trempus C, Mark W, et al. Deficiency of kinase suppressor of Ras1 prevents oncogenic ras signaling in mice. Cancer Res. 2003;63(14):4232–8.

70. Nguyen A, Burack WR, Stock JL, Kortum R, Chaika OV, Afkarian M, et al. Kinase suppressor of Ras (KSR) is a scaffold which facilitates mitogen-activated protein kinase activation in vivo. Mol Cell Biol. 2002;22(9):3035–45.

71. Kim YH, Lee JR, Hahn MJ. Regulation of inflammatory gene expression in macrophages by epithelial-stromal interaction 1 (Epsti1). Biochem Biophys Res Commun. 2018;496(2):778–83.

72. Roig AI, Eskiocak U, Hight SK, Kim SB, Delgado O, Souza RF, et al. Immortalized Epithelial Cells Derived From Human Colon Biopsies Express Stem Cell Markers and Differentiate In Vitro. Gastroenterology. 2010;138(3):1012-21.e5.

73. van de Wetering M, Francies HE, Francis JM, Bounova G, Iorio F, Pronk A, et al. Prospective derivation of a living organoid biobank of colorectal cancer patients. Cell. 2015;161(4):933–45.

74. Longo PA, Kavran JM, Kim MS, Leahy DJ. Transient mammalian cell transfection with polyethylenimine (PEI). Methods Enzymol. 2013;529:227–40.

75. Dobin A, Davis CA, Schlesinger F, Drenkow J, Zaleski C, Jha S, et al. STAR: ultrafast universal RNA-seq aligner. Bioinformatics. 2013;29(1):15–21.

76. Schmittgen TD, Livak KJ. Analyzing real-time PCR data by the comparative C(T) method. Nat Protoc. 2008;3(6):1101–8.

